# Interaction of YAP with the Myb-MuvB (MMB) complex defines a transcriptional program to promote the proliferation of cardiomyocytes

**DOI:** 10.1101/824755

**Authors:** Marco Gründl, Susanne Walz, Laura Hauf, Melissa Schwab, Kerstin Marcela Werner, Susanne Spahr, Carsten P. Ade, Stefan Gaubatz

**Author notes:** Correspondence to: Stefan Gaubatz.

## Abstract

The Hippo signalling pathway and its central effector YAP regulate proliferation of cardiomyocytes and growth of the heart. Using genetic models in mice we show that the increased proliferation of cardiomyocytes due to loss of the Hippo-signaling component SAV1 depends on the Myb-MuvB (MMB) complex. Similarly, proliferation of postnatal cardiomyocytes induced by constitutive active YAP requires MMB. Genome studies revealed that YAP and MMB regulate an overlapping set of cell cycle genes in cardiomyocytes. We find that YAP binds directly to B-MYB, a subunit of MMB, in a manner dependent on the YAP WW domains and a PPXY motif in B-MYB. Disruption of the interaction by overexpression of the YAP binding domain of B-MYB strongly inhibits the proliferation of cardiomyocytes. Our results point to MMB as a critical downstream effector of YAP in the control of cardiomyocyte proliferation.

## INTRODUCTION

The Hippo signaling pathway plays fundamental roles in proliferation and organ size control (*1*). In mammals, the Hippo cascade is composed of the MST1/2 and LATS1/2 kinases and the adaptor proteins MOB1 and Salvador (SAV1). When the Hippo pathway is active, LATS1/2 phosphorylate the transcriptional coactivators YAP and TAZ, which results either in their cytoplasmic retention by 14-3-3 proteins or SCF-mediated proteasomal degradation (*2*). In contrast, when Hippo is inactive, YAP/TAZ enter the nucleus and act as transcriptional coactivators predominantly by binding to DNA through TEAD transcription factors. The Hippo pathway and its downstream effectors YAP and TAZ are involved in cardiac development and have been implicated in heart regeneration after tissue damage (*3*, *4*). Active YAP promotes proliferation of postnatal cardiomyocytes and induces a fetal-like cell state in adult cardiomyocytes (*5*) (*6*). Similarly, cardiac-specific deletion of the Hippo kinases LATS1/2 or of the scaffold protein SAV1 results in heart enlargement and increased proliferation of embryonal and postnatal cardiomyocytes due to increased YAP activity (*7, 8*). Conversely, deletion of YAP suppresses cardiomyocyte proliferation (5, *7, 9*). Notably, activated YAP or loss of Hippo signaling can extend the neonatal proliferation of cardiomyocytes to postnatal stages, where proliferation is normally very low (*5*–*7*). In addition, recent studies report a better outcome after myocardial infarction in mice with hyperactivated YAP (*8*, *10-12*). Together, these studies identify YAP as an important regulator of cardiomyocyte proliferation and cardiac regeneration. However, the detailed mechanisms by which YAP promotes cardiomyocyte proliferation are still unclear (*13*).

We recently observed that Myb-MuvB complex (MMB) and YAP co-regulate an overlapping set of late cell cycle genes (*14*). MMB consist of the evolutionary conserved MuvB core of the five proteins LIN9, LIN52, LIN54, LIN37 and RBBP4 and associated proteins.

(*15*, *16*). Depending on its interactions with different binding partners, MuvB can either repress or activate transcription. Specifically, when MuvB interacts with the p130 retinoblastoma protein paralog, and with E2F4 and DP1, it forms the DREAM complex, which represses E2F-dependent gene expression in quiescent cells and in early G1 (*17–19*). Upon cell cycle entry, p130, E2F4 and DP1 dissociate from the complex and the MuvB core associates with the transcription factor B-MYB to form MMB (*19*, *20–22*). MMB mainly regulates genes required for mitosis and cytokinesis. Binding of YAP to enhancers promotes the activation of MMB-bound promoters, providing an explanation for the co-activation of late cell cycle genes by YAP and MMB (*14*).

Whether the ability of YAP to promote proliferation *in vivo* is mediated by MuvB complexes has not been addressed. To investigate the possible function of MuvB in YAP-mediated cardiomyocyte proliferation we used genetic approaches in mice and biochemical experiments. We find that YAP and MMB interact in the developing heart. *In vivo* in mice this interaction is essential for increased mitosis of Hippo-deficient cardiomyocytes. Additionally, we demonstrate that YAP driven proliferation of postmitotic cardiomyocytes is dependent on the interaction of YAP with the B-MYB subunit of MMB.

## RESULTS

### MuvB is required for proliferation and mitosis of Hippo-deficient embryonic cardiomyocytes

To explore the connection between the Hippo-YAP pathway and MuvB *in vivo* in the heart, we deleted the Hippo pathway member Salvador (*Sav1*), a scaffold protein that is required for Hippo kinase activity, and the MuvB subunit *Lin9* in cardiac precursor cells. Early cardiac specific deletion of *Lin9* and *Sav1* was achieved by using mouse strains with conditional (floxed) alleles of *Sav1* and *Lin9* in combination with Nkx2.5-Cre (*23*). Phosphorylation of YAP on S127 was reduced in Sav1 deficient hearts, indicating that YAP is hyperactivated (Figure S1). We first examined the effect of MMB inactivation on proliferation of *Sav1* mutated cardiomyocytes. To assess proliferation, we stained E13.5 heart sections for the cell cycle marker Ki67. Cardiomyocytes were identified by co-staining for cardiac troponin T (cTnT). The proportion of Ki67-positive cardiomyocytes was significantly increased in Nkx2.5-Cre;Sav^fl/fl^ hearts compared to control hearts with wildtype *Sav1*, indicating that loss of SAV1 promoted cardiomyocyte proliferation as has been reported previously (*7*) (Figure 1A,B). The fraction of cardiomyocytes staining positive for phosphorylated histone H3 (pH3), a marker of mitotic cells, was also increased in *Nkx2.5-Cre; Sav1*^*fl/fl*^ hearts (Figure 1C,D). In contrast, proliferation and mitosis were strongly reduced by inactivation of *Lin9* and the increase in proliferation by *Sav1* deletion was blocked in *Sav1, Lin9* double mutant cardiomyocytes (Figure 1A-D). These genetic experiments suggest that the proliferative phenotype due to the loss of the Hippo pathway member SAV1 is dependent on the MMB subunit LIN9.

**Figure 1:**
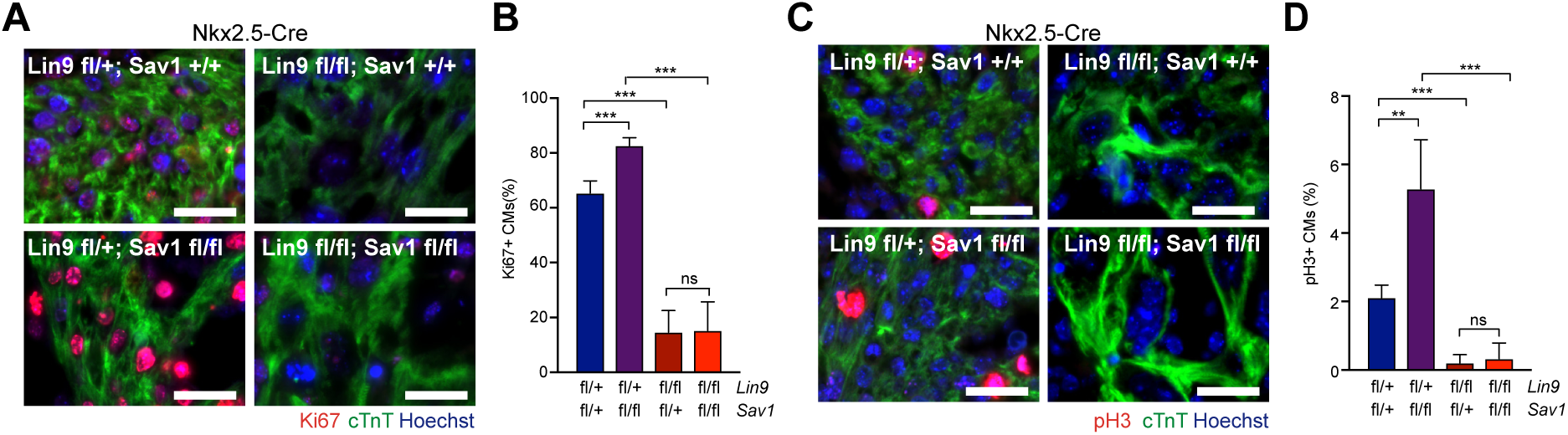
Proliferation of embryonic cardiomyocytes following Hippo inactivation depends on LIN9. A) and B) Heart sections from E13.5 mice with the indicated genotypes were stained for the proliferation marker Ki67 (red). Cardiomyocytes were identified by staining for cTnT (green). Example microphotographs are shown in (A), the quantification of Ki67-positive cells is shown in (B). Scale bar: 25µm. Error bars indicate SDs. Number of mice analyzed per genotype: *Lin9*^*fl/+*^;*Sav1*^*fl/+*^ n=5, *Lin9*^*fl/+*^; *Sav1*^*fl/fl*^ n=4, *Lin9*^*fl/fl*^; *Sav1*^*fl/+*^ n=6 and *Lin9*^*fl/fl*^;*Sav1*^*fl/fl*^ n=5 C) and D) Heart sections from E13.5 mice with the indicated genotypes were stained for the mitosis marker pH3 (red). Cardiomyocytes were identified by staining for cTnT (green). Example microphotographs are shown in (C), the quantification of pH3-positive cells is shown in (D). Number of mice analyzed per genotype: *Lin9*^*fl/+*^;*Sav1*^*fl/+*^ n=5, *Lin9*^*fl/+*^; *Sav1*^*fl/fl*^ n=4, *Lin9*^*fl/fl*^; *Sav1*^*fl/+*^ n=6 and *Lin9*^*fl/fl*^;*Sav1*^*fl/fl*^ n=6. Error bars indicate SD. B, D: Student’s t-test. *=p<0.05, **=p<0.01, ***=p<0.001, ns= not significant.

### Hippo and MMB regulate an overlapping set of genes in embryonic cardiomyocytes

Next, we explored the impact of *Lin9* mutation on the transcriptional program of Hippo-deficient cardiomyocytes by performing RNA-seq using RNA isolated from embryonic heart ventricles of mice carrying cardiac specific deletions of *Sav1* alone, *Lin9* alone or *Sav1* and *Lin9* together. In *Sav1* knockout hearts, E2F target genes and other gene sets related to cell cycle regulation were enriched after inactivation of Hippo-signaling (Figure 2A, Supplementary Figure S2A). Strikingly, the activation of these gene sets was suppressed by the simultaneous deletion of *Lin9* together with *Sav1*, indicating that the elevated expression of these genes in *Sav1* mutant hearts requires the function of MMB (Figure 2A, B, Supplementary Figure S2A). By qPCR we independently validated that a set of cell cycle target genes were activated in *Sav1* mutated hearts, downregulated in *Lin9* mutated hearts and remained downregulated in hearts isolated from double mutant embryos (Figure 2C).

**Figure 2:**
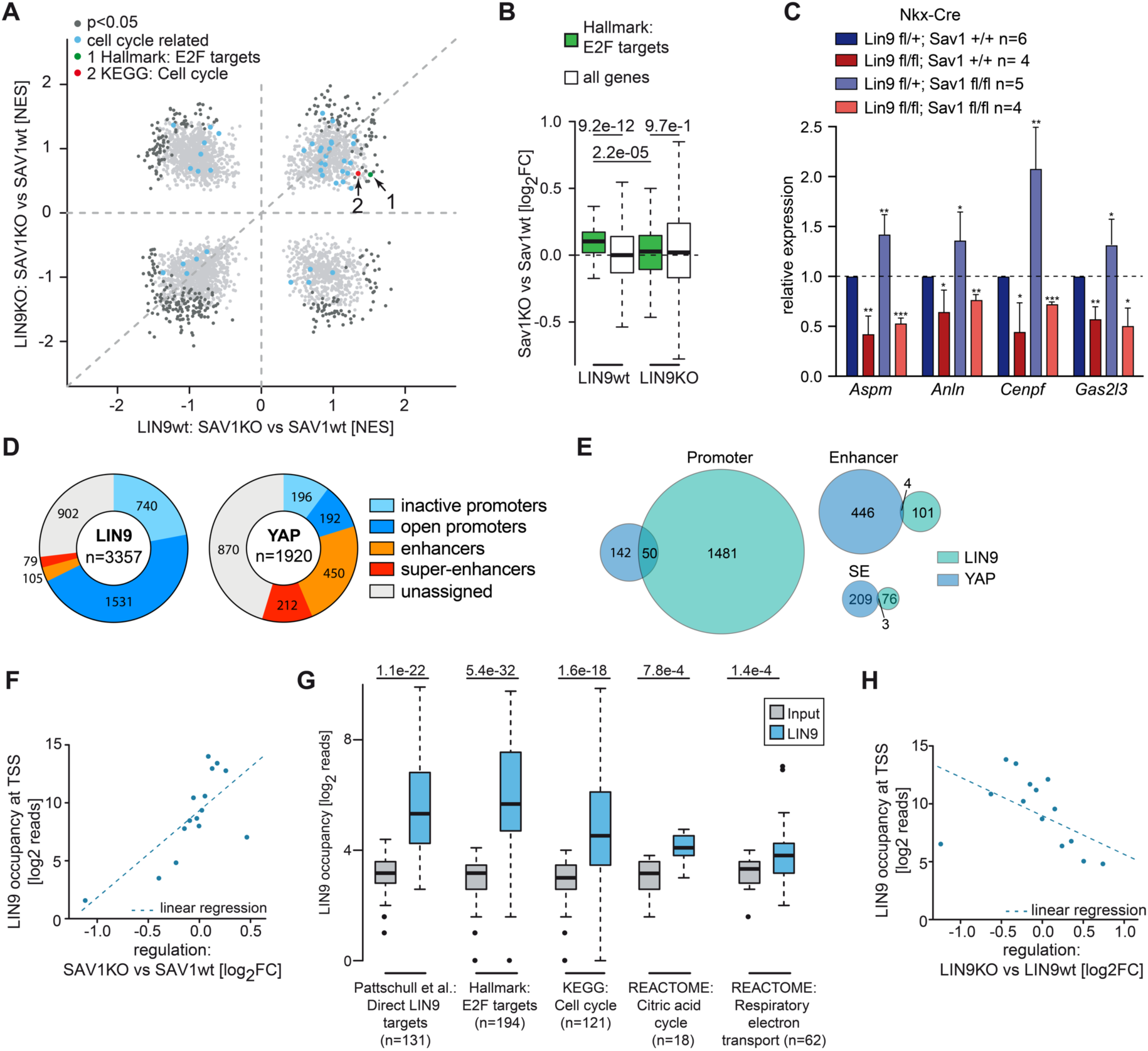
Cell cycle genes upregulated in *Sav1* knockout hearts are direct targets of LIN9. A) GSEA comparing the effect of deletion of Sav1 in Nkx-Cre LIN9 wt and LIN9 KO heart ventricles at E13.5. Gene sets related to cell cycle are highlighted in blue. NES: normalized enrichment score. B) Boxplot comparing differences in E2F target gene expression between Nkx-Cre; Sav1^fl/fl^ (Sav1 KO) and Nkx-Cre; Sav1^+/+^ (Sav1 wt) heart ventricles in LIN9^fl/fl^ (LIN9 KO) or Lin9^fl/+^ (LIN9 wt) background. C) Expression of the indicated genes relative to actin and *Hprt* in heart ventricles of embryos with the indicated genotypes was analyzed by RT-qPCR. D) Genomic localization of LIN9 and YAP in fetal (E16.5) heart ventricles. Active promoters, enhancers and super-enhancers are defined based on publicly available ChIP-seq data for histone marks. E) Venn diagram showing the overlap between binding by LIN9 and YAP in E16.5 hearts. F) Bin plot correlating changes in gene expression in Nkx-Cre; Lin9 ^fl/fl^ heart ventricles with binding of LIN9 to the promoter. Analyzed was a region 1kb upstream the TSS and input signals were subtracted. 15,642 expressed genes were grouped into 15 bins and the mean of each bin is plotted. Dashed line: Regression based on a linear model. G) Boxplot depicting LIN9 binding to genes from the gene sets shown in Supplementary Figure S2A and S2D. Analyzed was a region of +/-250bp around the TSS. H) Bin plot correlating changes in gene expression after *Sav1* knockout in Nkx-Cre hearts with binding of LIN9 to the promoters at E16.5. Analysis was performed as in F.

To find out whether cell cycle genes upregulated in *Sav1* hearts are direct targets of MMB, we performed chromatin immunoprecipitation experiments followed by high-throughput sequencing (ChIP-seq). We identified 3357 bindings sites for LIN9 in the fetal heart. By comparison with previously reported ChIP-seq data of histone modifications characteristic for active promoters and enhancers, most LIN9 binding sites are located in active promoters while less than 5% of LIN9 sites are found in enhancers or super-enhancers (Figure 2D, Supplementary Figure S2B). ChIP-seq of YAP in E16.5 hearts showed that YAP predominantly binds to enhancers and super-enhancers rather than to promoters (Figure 2D), which is consistent with recent genome wide studies in human cell lines (*14*, *24*–*26*). Consequently, there was little overlap in the binding of YAP and LIN9 (Figure 2E, Supplementary Figure S2B). To identify the direct targets of LIN9 in the heart, we plotted changes in gene expression upon deletion of *Lin9* against the density of promoter-bound LIN9. This revealed a correlation between genes that are activated by LIN9 (i.e. that are downregulated after deletion of *Lin9*) and promoter binding of LIN9 (Figure 2F). In particular, LIN9 strongly bound to the promoters of cell cycle genes and E2F target genes (Figure 2G). While overall LIN9-binding correlated with expression changes, LIN9 only weakly bound to the promoters of some gene sets that are also downregulated in *Lin9* knockout hearts, including genes related to mitochondrial function, oxidative phosphorylation, metabolism, heart muscle contraction and ion channel activity (Figure 2G, Supplementary Figure S2C,D). Downregulation of these genes is likely an indirect consequence of loss of *Lin9*. Strikingly, plotting changes in gene expression upon cardiac-specific deletion of *Sav1* against LIN9-promoter occupancy showed that promoters of genes activated in *Sav1* knockout hearts were bound by LIN9, indicating that they are direct targets of MMB (Figure 2H). Altogether these data support the idea that YAP activates LIN9-bound cell cycle promoters from distant enhancers as previously shown in human MCF10A cells (*14*). In summary, the deletion of *Sav1* hyperactivates YAP, which in turn results in induction of cell cycle genes in a LIN9 dependent manner.

### Lin9 is required for proliferation of Hippo-deficient postnatal cardiomyocytes

Deletion of *Lin9* in heart progenitor cells by Nkx2.5-Cre resulted in embryonic lethality due to defects in cardiomyocyte proliferation and division resulting in enlarged nuclei and leading to reduced thickness of the compact myocardium of both ventricles (Figure 3A-C, Supplementary Figure 3A). The phenotype of Nkx2.5-Cre; *Lin9*^*fl/f*l^ mice is consistent with the reduced expression of cell cycle genes in the heart. To circumvent the embryonic lethality associated with deleting *Lin9* in early cardiac precursors, we used a transgenic mouse line with Cre recombinase driven by the alpha-MHC-promoter, which is active at a later stage during heart development compared to Nkx2.5-Cre (*27*). α-MHC-Cre; *Lin9*^*fl/fl*^ mice survived into adulthood without any obvious heart phenotype and without differences in the heart to body weight when compared to heterozygous control animals (Figure 3D-F, Supplementary Figure 3B). The lack of a heart phenotype in α-MHC-Cre; *Lin9*^*fl/fl*^ mice is not due to inefficient deletion of *Lin9* in the heart as revealed by combining with a mT/mG reporter strain (*28*). Upon Cre-induced recombination of the mT/mG reporter gene, membrane bound Tomato (mT) is removed and the expression of membrane bound EGFP (mG) is activated (Figure 3G). FACS analysis revealed that recombination in the heart of α-MHC-Cre; *Lin9*^*fl/fl*^ mice was almost 100% (Figure 3G, Supplementary Figure 3C).

**Figure 3:**
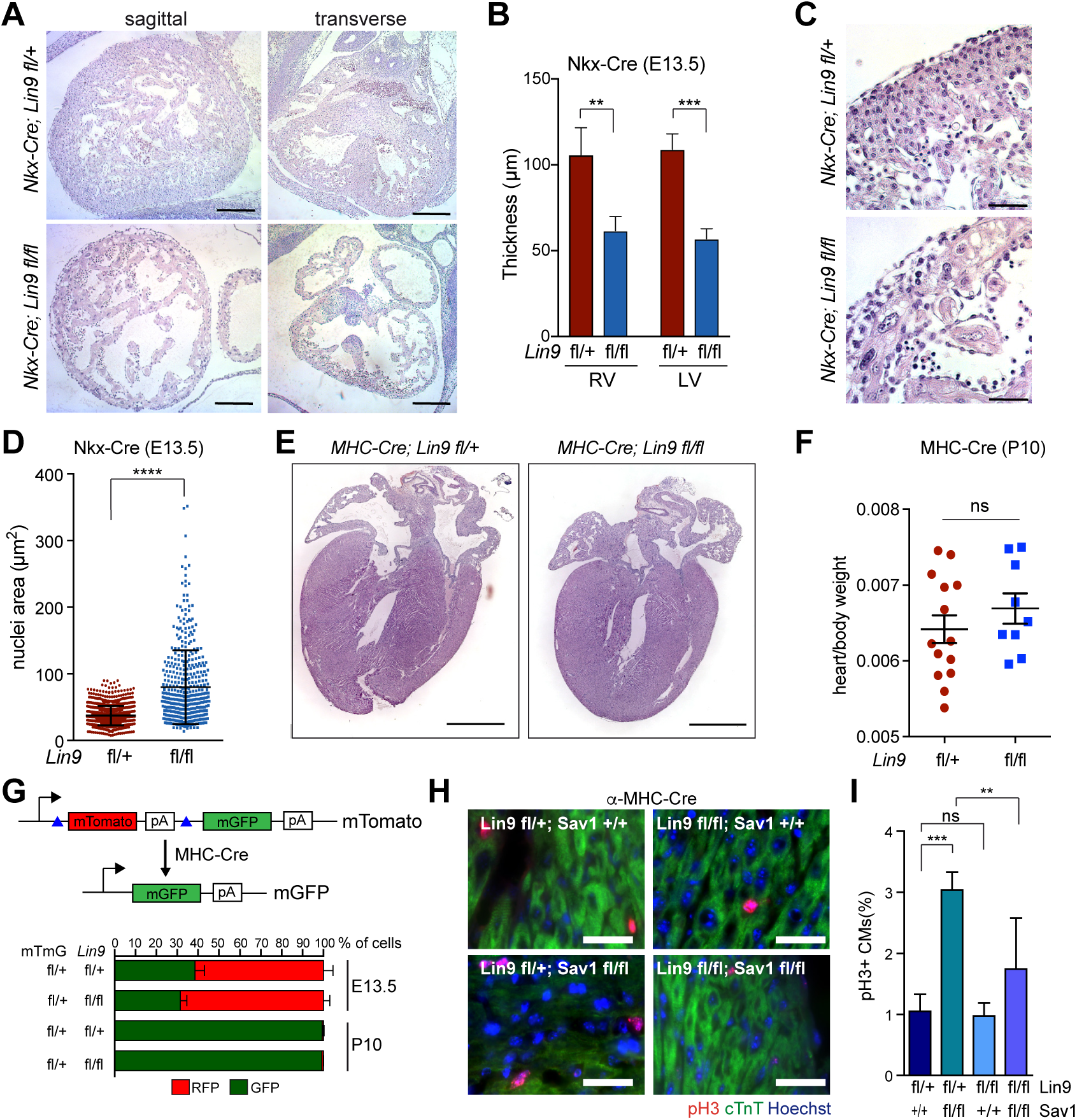
LIN9 is required for cardiomyocyte proliferation in Hippo-deficient postnatal hearts. A) H&E-stained sagittal and transverse sections of embryonic E13.5 hearts of mice with the indicated genotypes. Scale bar: 100µm B) Thickness of the right and left ventricular compact myocardium. n=5 *Lin9*^*fl/+*^ hearts and n=3 *Lin9*^*fl/fl*^ hearts. C) High magnification of the myocardium of *Lin9*^*fl/fl*^ embryonic hearts demonstrating pleomorphic and enlarged nuclei. Scale bar: 25µm D) Quantification of the nuclear area in the myocardium of *Lin9*^*fl/+*^ and *Lin9*^*fl/fl*^ embryonic hearts. E) H&E-stained sections of hearts from α-MHC-Cre*;Lin9*^*fl/+*^ and α-MHC-Cre;*Lin9*^*fl/fl*^ mice at P10. Scale bar: 1 mm F) heart to body weight of α-MHC-Cre; *Lin9*^*fl/+*^ and α-MHC-Cre;*Lin9*^*fl/fl*^ mice at p10. n= 14 control mice and n=9 *Lin9*^*fl/fl*^ mice. G) Scheme of the mT/mG reporter. Upon expression of Cre-recombinase, mTomato is deleted and mGFP is activated. Quantification of mTomato (mT) and mEGFP (mG) positive cardiomyoyctes. E13.5 *Lin9*^*fl/+*^ n=4; E13.5 *Lin9*^*fl/fl*^ n=4; P10 *Lin9*^*fl/+*^ n=3; P10 *Lin9*^*fl/fl*^ n=3. H) and I) Fraction of pH3-positive cardiomyocytes in the hearts of 10 days old mice with the indicated genotypes. Cardiomyocytes were identified by staining for cTnT. Scale bar 25µm. *Lin9*^*fl/+*^; *Sav1*^*fl/+*^ n=3, *Lin9*^*fl/+*^; *Sav1*^*fl/fl*^ n=6, *Lin9*^*fl/fl*^; *Sav1*^*fl/+*^ n=3 and *Lin9f*^*l/fl*^;*Sav1*^*fl/fl*^ n=7. B),D), I): Error bars indicate SDs. Error bars indicate SEM. Student’s t-test. *=p<0.05, **=p<0.01, ***=p<0.001, ns= not significant.

Also, LIN9 was expressed and remained associated with chromatin in postnatal hearts as determined by ChIP seq (Supplementary Figure S3D,E,F). The overall chromatin binding pattern of LIN9 was similar in embryonic and postnatal hearts and the shared binding sites reflect high confident LIN9-binding sites (Supplementary Figure S3F,G,H).

To address whether LIN9 is required for mitosis of postnatal cardiomyocytes in Hippo-deficient hearts, we next crossed α-*MHC-Cre; Lin9*^*fl/fl*^ mice to *Sav1*^*fl/fl*^ mice. Under normal conditions, there are almost no dividing cardiomyocytes in the postnatal heart, as expected. The deletion of *Sav1* robustly increased the fraction of mitotic cardiomyocytes (Figure 3H,I). Strikingly, this phenotype was suppressed when *Lin9* was lost and the fraction of mitotic cardiomyocytes remained low in α-MHC-Cre; *Sav1*^*fl/fl*^, *Lin9*^*fl/fl*^ double mutant mice, indicating a role for MMB in mitotic entry of postnatal cardiomyocytes due to Hippo deficiency (Figure 3H,I).

Thus, LIN9 is dispensable in the postnatal heart probably because cardiomyocytes hardly divide at this time point. However, LIN9 becomes necessary for ectopic cardiomyocyte proliferation in the absence of SAV1.

### Cardiomyocyte proliferation by activated YAP requires LIN9

To directly test whether increased cardiomyocyte proliferation due to activated YAP depends on MMB, we transduced cardiomyocytes with an adenovirus expressing a constitutive active version of YAP in which S127, the site of the inactivating phosphorylation by LATS kinases, is mutated to alanine (Figure 4A). To determine the requirement of MMB for YAP(S127A) induced proliferation, we used cardiomyocytes isolated from mice with a conditional allele of *Lin9* (*Lin9*^*fl/fl*^) and expressing a hormone inducible CreER recombinase that can be activated by the addition of 4-hydroxytamoxifen (4-OHT). Treatment with 4-OHT resulted in efficient deletion of *Lin9 in* neonatal *Lin9*^*fl/fl*^; CreER cardiomyocytes (Figure 4B). Expression of YAP(S127A) robustly induced mitotic entry of embryonic E14.5 and postnatal P1 cardiomyocytes, as reported before (*5*, *9*) (Figure 4C,D, Supplementary Figure S4A,B). Importantly, when *Lin9* was deleted by treatment with 4-OHT, the increase in pH3 positive cells by YAP(S127A) was strongly suppressed, indicating that YAP requires MMB to promote cardiomyocyte proliferation (Figure 4C,D, Supplementary Figure S4A,B).

**Figure 4:**
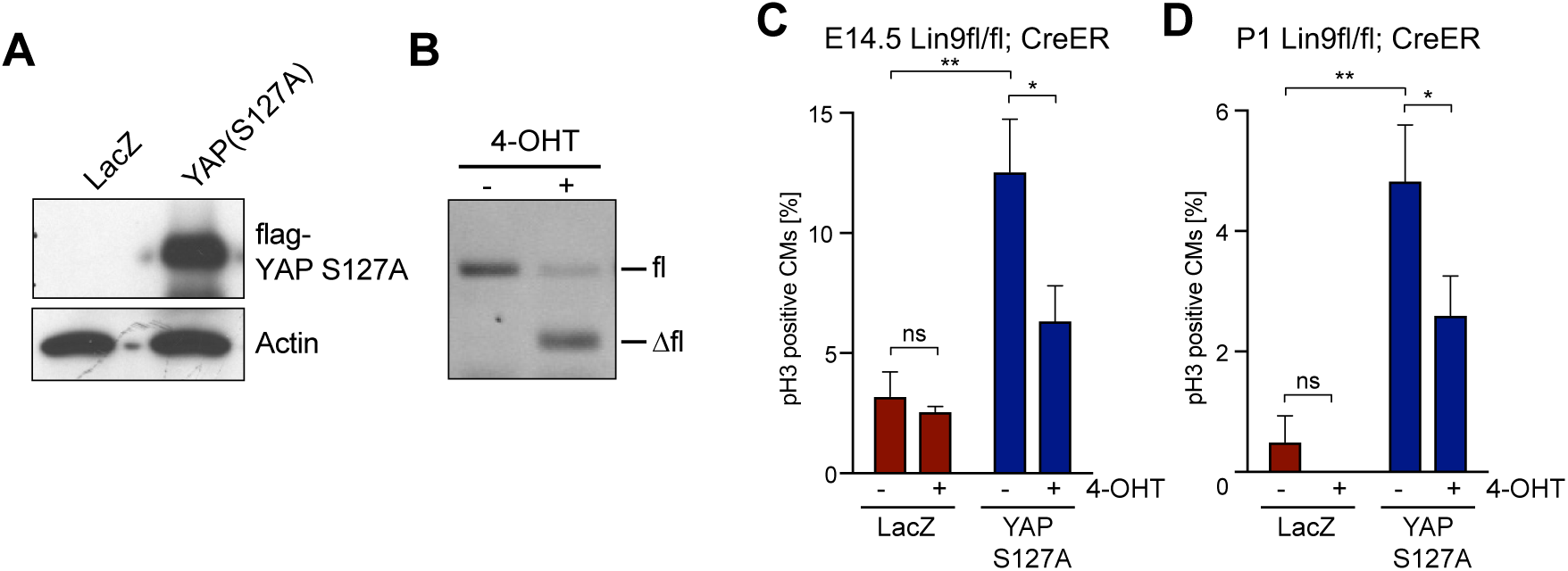
Cardiomyocyte proliferation by activated YAP requires MMB. A) Cardiomyocytes isolated from neonatal (P1) *Lin9*^*fl/fl*^;CreER mice were transduced with a LacZ control (Ade-LacZ) or FLAG-YAP(S127A) expressing adenovirus. Expression of YAPS127A was verified by immunoblotting with a flag-antibody 72 hours after adenoviral transduction. Actin served as a control. B) Cardiomyocytes isolated from *Lin9*^*fl/fl*^;CreER hearts were treated without and with 4-OHT as indicated and the deletion of *Lin9* was verified by genomic PCR. C) and D) Embryonal (E14.5) *Lin9*^*fl/fl*^;CreER cardiomyocytes (C) or postnatal (P1) cardiomyocytes (D) were transduced with Ade-LacZ or with Ade-YAP(S127A) and treated with or without 4-OHT. The fraction of pH3-positive cardiomyocytes was quantified. Example microphotographs are shown in Supplementary Figure S4. Error bars show SDs of biological replicates (n=3).

### YAP physically interacts with MMB in cardiomyocytes

YAP and MMB could independently regulate a similar set of genes required for cell cycle regulation or they could cooperate by binding to each other. To address these possibilities, we next investigated whether YAP and B-MYB interact in the heart. Proximity ligation assays (PLA) showed that YAP indeed interacted with B-MYB in developing cardiomyocytes (Figure 5A,B). The interaction was specific, as demonstrated by the loss of the PLA signal upon siRNA-mediated depletion of B-MYB (Figure 5A,B). YAP also interacted with LIN9, a core subunit of the MuvB complex that is required for binding of B-MYB to MuvB (*29*), providing further evidence for an interaction between YAP and MMB in cardiomyocytes.

**Figure 5:**
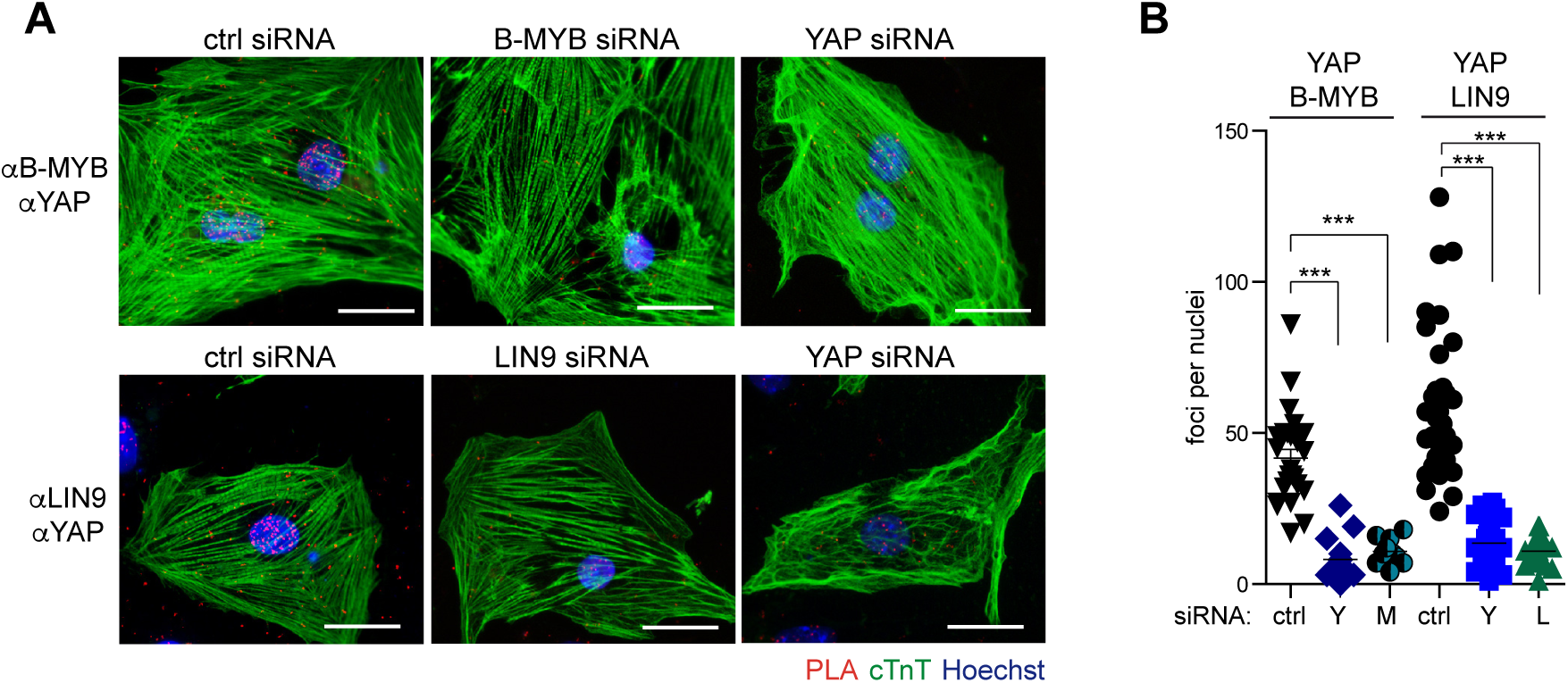
Interaction between YAP and MMB in cardiomyocytes. A) Proximity ligation assays (PLA) showing that YAP and LIN9 and YAP and B-MYB interact in the nuclei of E14.5 cardiomyocytes. siRNA mediated depletion of YAP/TAZ, LIN9 or B-MYB served as control. Interactions are indicated by red fluorescent dots. Cardiomyocytes were identified by immunostaining for cTnT (green). Bar: 25 µm B) PLA assays shown in A were quantified by counting the number of interactions per nucleus.

### B-MYB interacts with the tandem WW domains of YAP

To gain further insights into the interplay between YAP and B-MYB and to develop tools to interfere with the interaction, we mapped the domains of YAP and B-MYB that are involved in the interaction. In co-immunoprecipitation experiments with HA-B-MYB and a set of truncated flag-tagged YAP constructs, B-MYB interacted with YAP only when the tandem WW domains were present. B-MYB did not interact with the C-terminal transactivation domain or the PDZ-binding motif (Figure 6A,B, Supplementary Figure S5A). Internal deletion of the tandem WW domains abolished the binding, confirming that these domains are required for the YAP-B-MYB interaction (Figure 6C). To verify that the YAP WW domains mediate the interaction with B-MYB, we performed pulldown experiments with recombinant GST fused to the N-terminal part of YAP containing the TEAD-binding and WW domains (GST-TEAD-WW1/2) or just the two WW domains (GST-WW1/2) (Supplementary Figure S5B). HA-B-MYB specifically bound to GST-TEAD-WW1/2 and GST-WW1/2 and but not to GST alone (Figure 6D). As a control, HA-TEAD4 only bound to GST-TEAD-WW1/2 containing the TEAD-binding domain, but not to GST alone or to GST-WW1/2. HA-tagged EB1, which was used as a negative control, did not bind to any of the GST constructs. Although B-MYB can independently bind to both WW domains, it more strongly interacted with the first WW domain and strongest binding of B-MYB was observed when both WW domains were present (Supplementary Figure S5C).

**Figure 6:**
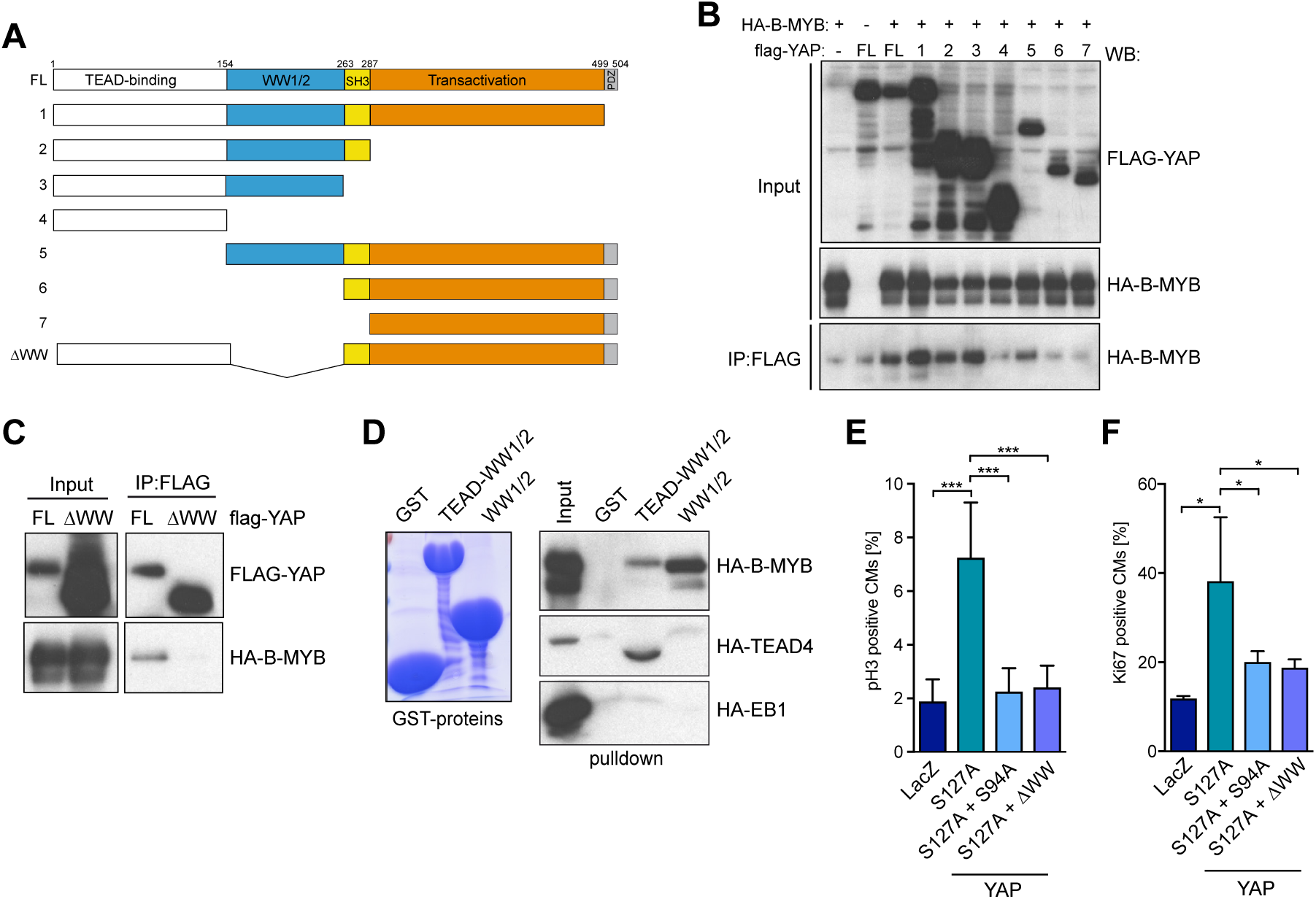
The WW domains of YAP mediate the interaction with B-MYB and are required to induce cardiomyocyte proliferation. A) Scheme of YAP deletion mutants used in co-immunoprecipitation experiments. B) Co-immunoprecipitation of ectopically expressed HA-B-MYB with flag-tagged YAP. Lysates of HeLa cells expressing HA-B-MYB and flag-YAP mutants (see A) were immunoprecipitated with flag-antiserum and immunoblotted with an anti-HA-antibody. 3 % percent of total lysate was immunoblotted (Input). A quantification is provided in Supplementary Figure S5A. C) Co-immunoprecipitation between flag-YAP and HA-B-MYB demonstrating that the WW domains of YAP are required for the interaction with B-MYB. D) Pulldown experiments with 5 µg of the indicated recombinant GST-fusion proteins and with HA-B-MYB, HA-TEAD4 and HA-EB1 expressed in HeLa cells. Bound proteins were detected with an anti-HA antibody. HA-TEAD was used as a positive control for the interaction with GST-YAP-TEAD-WW1/2 and HA-EB1 served as a negative control. A coomassie brilliant blue stained gel of the purified proteins is shown to the left. E) and F) Cardiomyocytes were transduced with Ade-LacZ, Ade-YAP(S127A), and Ade-YAP(S127A/S94A) or with Ade-YAP(S127A/∆WW). The fraction of pH3-positive positive (E) and Ki67 (F) cardiomyocytes was determined. Example microphotographs are shown in Supplementary Figure S6E and F. Error bars show SD. (E) n=6-8, (F) n=3-4 biological replicates.

Importantly, the ability of YAP(S127A) to stimulate cardiomyocyte proliferation and to promote entry into mitosis was abolished by deletion of the YAP WW domains, which mediate the binding to MMB (Figure 6E,F, Supplementary Figure S5D,E). Similarly, a point mutant of YAPS127A deficient in binding to TEAD [YAP(S127A/S94A)], was also not able to increase proliferation and mitosis, indicating that these functions of YAP are TEAD dependent.

### Overexpression of the YAP binding domain disrupts the B-MYB-YAP interaction and inhibits proliferation

Since YAP binds to the N-terminus of B-MYB and since the MMB-interaction domain (MBD) is located in the C-terminal region, the binding of YAP to B-MYB is likely not mediated by binding of YAP to the MuvB core (*14*, *29*). In pulldown experiments with GST-WW1 and with a set of HA-tagged B-MYB deletion mutants we mapped the minimal binding site for YAP to amino acids 80 and 241 of B-MYB (Figure 7A,B). For example, B-MYB(2-241) robustly bound to YAP whereas B-MYB(2-79) and B-MYB(242-410) failed to bind to YAP (Figure 7B). The interaction between B-MYB and YAP could be indirect since exogenously expressed HA-B-MYB likely associates with additional cellular proteins. To test whether the interaction between B-MYB and YAP is direct, we incubated purified, recombinant his-tagged B-MYB(2-241) with GST-WW1/2. Purified his-B-MYB(2-241) interacted with GST-WW1/2, indicating that the binding between B-MYB and YAP is direct (Figure 7C).

**Figure 7:**
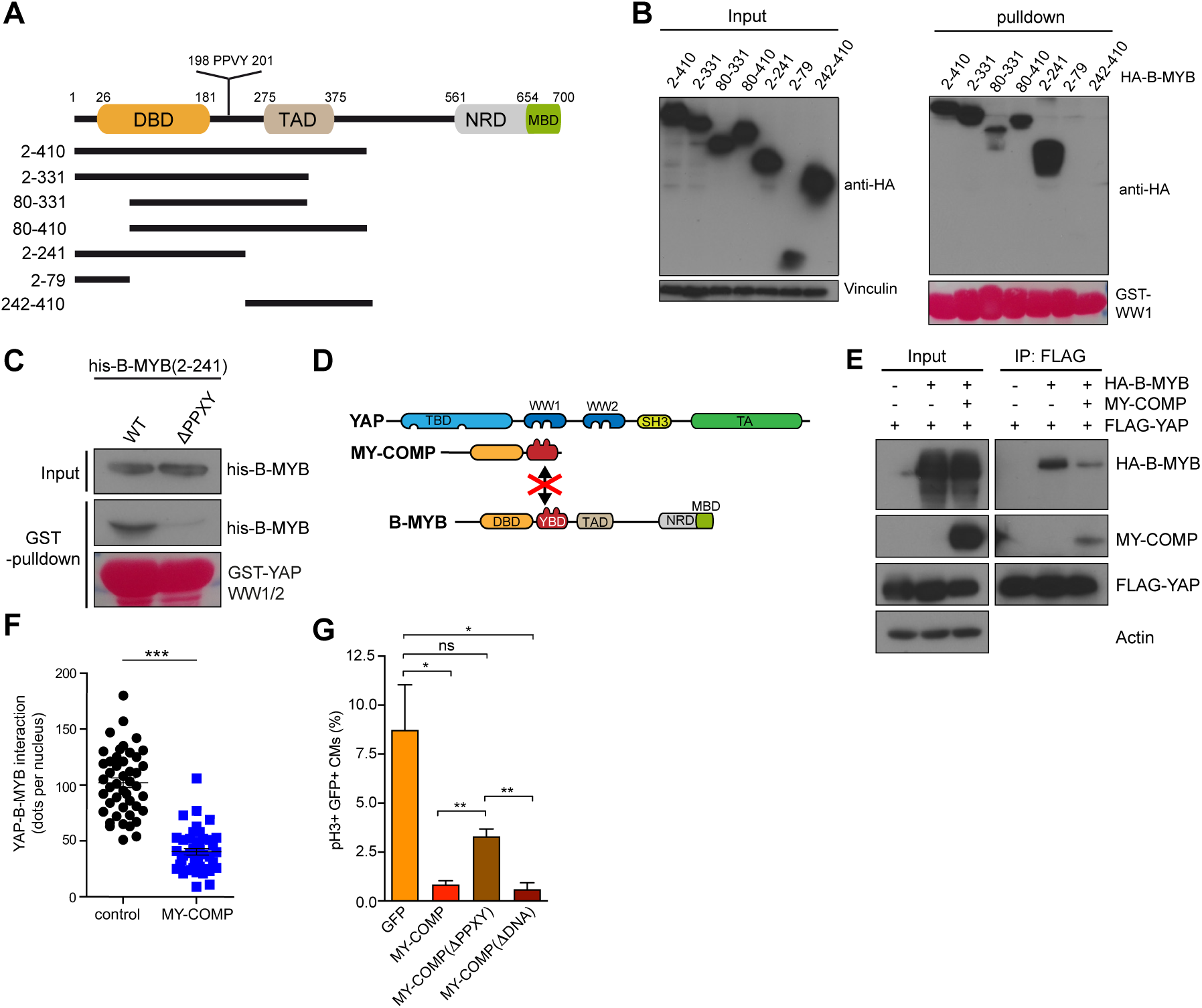
Disrupting the association between B-MYB and YAP by MY-COMP inhibits cardiomyocyte proliferation. A) Scheme of the domain structure of B-MYB and deletion constructs. B) Pulldown experiments with GST-WW1 and HA-tagged B-MYB constructs. Bound proteins were detected by immunoblotting with a HA antibody. Input: 3 % of the lysate. Vinculin was used as a control. Recombinant GST-proteins were detected by Ponceau staining. C) Pulldown experiments with GST-WW and his-tagged B-MYB(2-241) and B-MYB(2-241,∆PPXY). Bound B-MYB was detected with a his-antibody. Input: 3 % of the recombinant his-tagged B-MYB constructs. Recombinant proteins were detected by Ponceau S staining. D) Scheme for the disruption of the YAP-B-MYB interaction by MY-COMP. E) HeLa cells were transfected with flag-YAP, HA-B-MYB and MY-COMP (HA-NLS-B-MYB(2-241)) as indicated. Flag-YAP was immunoprecipitation and bound HA-B-MYB was detected by immunoblotting. When MY-COMP was co-transfected, the interaction between flag-YAP and HA-B-MYB was decreased. Input: 3 % of the lysate. A quantification is shown in Suppl. Fig. S6C. F) PLA of endogenous YAP and B-MYB upon transfection of MY-COMP showing that the MY-COMP disrupts the interaction between YAP and B-MYB. Example microphotographs are shown in Suppl. Fig. S6D. G) Disruption of the YAP-B-MYB interaction by MY-COMP inhibits mitosis of cardiomyocytes. Embryonal cardiomyocytes were infected with adenoviruses expressing GFP, MY-COMP, MY-COMP,∆PPXY or with MYCOMP,∆DNA each coupled to GFP through a T2A self-cleaving peptide. Infected cardiomyocytes were detected by staining for cTnT and by their green fluorescence. Mitotic cells were quantified by staining for phospho-H3. Example microphotographs are provided in Suppl. Fig. S6. F), G) Error bars indicate SEMs. Student’s t-test. *=p<0.05, **=p<0.01, ***=p<0.001, ns= not significant.

The WW domain is a protein-interaction domain characterized by two tryptophan-residues separated by 20 to 22 amino acids. Ligands for the WW domain are proline-rich regions such as the PPXY motif (*30*, *31*). B-MYB harbors a PPXY sequence (PPVY) between the N-terminal DNA-binding domain and the central transactivation domain (Figure 7A). Importantly, this PPXY motif is located in the region of B-MYB that was found to interact with YAP, leading to the hypothesis that it is involved in binding to YAP. We directly tested this idea by performing pulldown experiments with a deletion mutant of recombinant B-MYB lacking the PPXY sequence (∆PPXY). Compared to his-tagged B-MYB(2-241), binding of his-B-MYB(2-241,∆PPXY) to the WW domains of YAP was strongly reduced (Figure 7C). The PPXY motif was also required for the interaction with YAP in the context of the full-length B-MYB protein in cells (Supplementary Figure S6A,B). Taken together, these results indicate that the YAP interacting region of B-MYB is located between amino acid 80 and 241 of B-MYB and involves a PPXY motif.

Since B-MYB and YAP directly interact, we next asked whether overexpression of the YAP binding domain of B-MYB can interfere with the interaction of B-MYB with YAP due to steric hindrance (Figure 7D). To address this possibility we created B-MYB(2-241) fused to a HA tag and a nuclear localization signal (HA-NLS-B-MYB-2-241), and named it MY-COMP for <u>M</u>YB-<u>Y</u>AP <u>comp</u>etition. First, we expressed MY-COMP in cells and performed co-immunoprecipitation experiments. In HeLa cells that express MY-COMP, the amount of full length HA-B-MYB co-precipitating with flag-YAP was strongly reduced when compared to control transfected cells (Figure 7E, Supplementary Figure S6C). Expression of MY-COMP also interfered with the endogenous YAP and B-MYB interaction as determined by proximity ligation assays (PLA) (Figure 7F, Supplementary Figure S6D).

We next infected embryonal cardiomyocytes with an adenovirus expressing MY-COMP coupled to GFP through a T2A self-cleaving peptide. Infected cardiomyocytes were detected by their green fluorescence and by staining for cTnT (Supplementary Figure S6E). Strikingly, staining for phospho-H3 showed that expression of MY-COMP strongly suppressed mitosis of embryonal cardiomyocytes (Figure 7G, Supplementary Figure S6E). Importantly, this effect was diminished by deletion of the PPXY motif in MY-COMP, indicating that the ability to prevent proliferation correlates with the ability to disrupt the YAP-B-MYB interaction. The effect of MY-COMP is not due to interference with the DNA-binding of B-MYB, because expression of MY-COMP with a mutation in the DNA-binding domain that has been shown to prevent the interaction with DNA (*32*), showed the same phenotype. Taken together these observations are consistent with the notion that the YAP-induced cardiomyocyte proliferation is mediated by the interaction of YAP with MMB.

### MMB target genes are downregulated in differentiated cells and re-activated by YAP

In another line of evidence, we investigated whether YAP is able to regulate MMB target genes in differentiated C2C12 myotubes. Co-immunoprecipitations confirmed that B-MYB and YAP interact in C2C12 myoblasts (Figure 8A). MMB target genes were expressed at high levels in asynchronous growing C2C12 cells and were downregulated during myogenic differentiation (Figure 8B). Expression of a constitutive active YAP(S127A) partially reverted the downregulation of MMB target genes in differentiated cells (Figure 8C). Importantly, like the induction of proliferation of cardiomyocytes, the upregulation of MMB target genes by YAP was dependent on the WW and the TEAD-binding domains (Figure 8D). Strikingly activation of YAP target genes that are not regulated from distant enhancers but by binding of YAP to the proximal promoter, such as *Ctgf* and *Cyr61* did not require the WW domains, indicating that it is independent from binding to MMB. Similar to what was observed in C2C12 cells, MMB target genes such as *Cenpf*, *Nusap1*, *Top2a* and *Birc5* were expressed at high levels in embryonic hearts when cardiomyocytes still divide but strongly declined in P1 and P10 hearts when cardiomyocytes differentiate and exit the cycle (Figure 8E,F). *Mybl2*, also declined at P1 compared to E16.5 and further decreased at P10, a pattern that was also confirmed on protein level (Figure 8E,F). Thus downregulation of MMB target genes correlates with the postnatal cell cycle exit of cardiomyocytes. Conversely, *Mybl2* and mitotic MMB target genes were re-activated in P1 *Sav1* knockout hearts when YAP is activated (Figure 8G). Together these data the support the view that MMB contributes to the induction of cell cycle genes in response to Hippo-deficiency or YAP activation.

**Figure 8:**
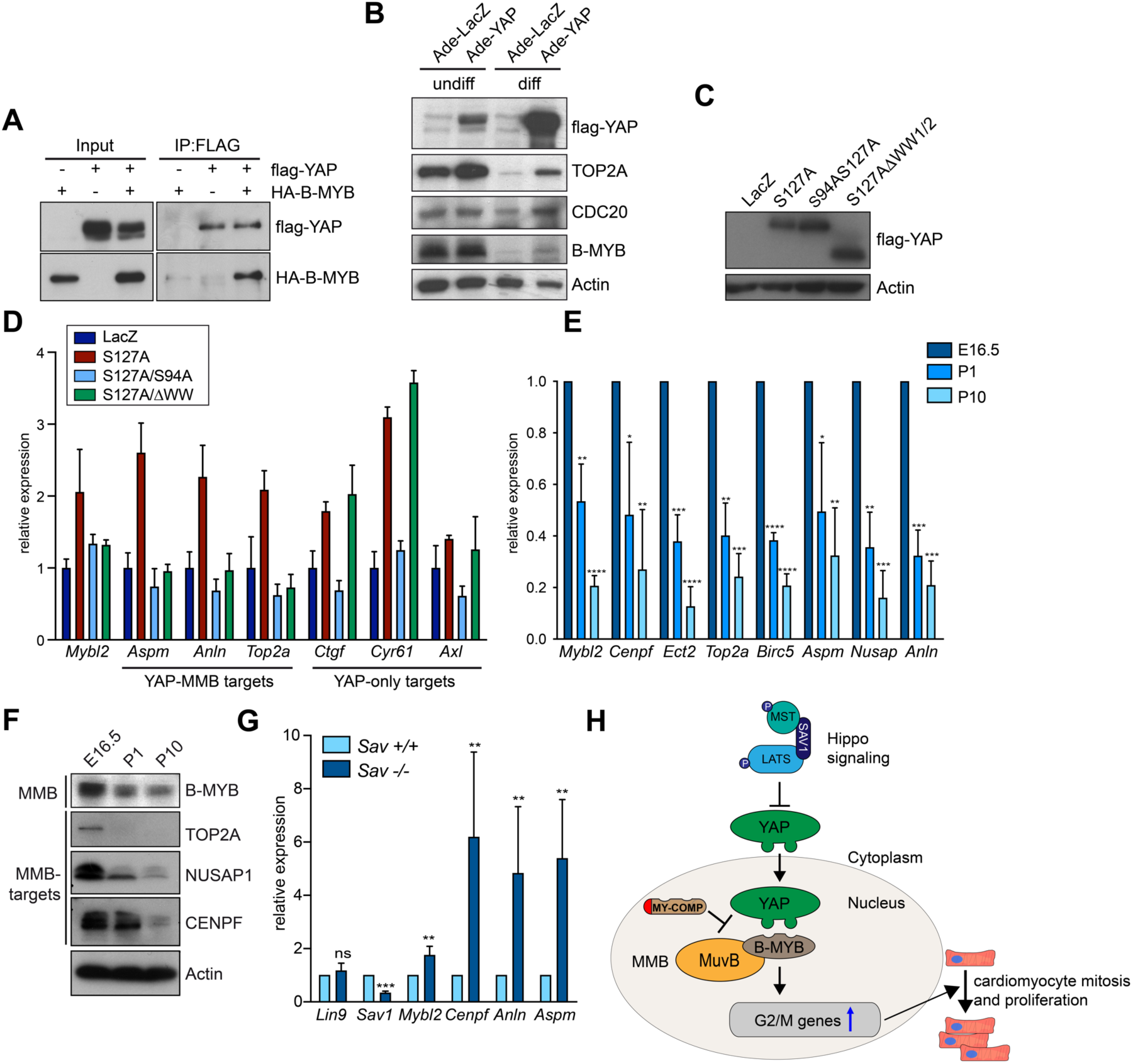
MMB target genes are downregulated in differentiated cells and re-activated by YAP. A) Co-immunoprecipitation of ectopically expressed flag-YAP and HA-B-MYB in C2C12 cells. IP, immunoprecipitation. Input: 3% of the amount used for IP was used. B) Undifferentiated C2C12 myoblasts (undiff) or differentiated C2C12 myotubes (diff) were infected with Ade-LacZ or Ade-YAP(S127A). After 2 days, the expression of the indicated proteins was analyzed by immunoblotting. Actin served as a loading control. C) and D) Differentiated C2C12 myotubes were infected with the indicated adenoviral expression constructs. C) The expression of the YAP constructs was analyzed by immunoblotting with a flag-antibody. Actin served as a loading control. D) mRNA expression of the indicated genes relative to *Actin* and *Hprt* was analyzed by RT-qPCR. Error bars show SD of technical triplicates from one representative experiment (n=3). E) The expression of the indicated MMB target genes relative to *Hprt and Actin* was investigated in E16.5, P1 and P10 hearts. n= 3 independent replicates. F) The expression of the indicated proteins in lysates prepared from hearts at the different developmental stages was investigated by immunoblotting. β-Actin served as a control. G) Expression of the indicated genes in Nkx2.5-Cre;*Sav1*^*+/+*^ and Nkx2.5-Cre;*Sav1*^*fl/fl*^ in hearts of P1 mice was investigated. n= 7 independent Nkx2.5-Cre;*Sav1*^*+/+*^ and Nkx2.5-Cre;*Sav1*^*fl/fl*^ animals. H) Model summarizing the results. See text for details. E), G) Error bars indicate SD. Student’s t-test. *=p<0.05, **=p<0.01, ***=p<0.001, ns= not significant.

## DISCUSSION

Recent studies have shown that the Hippo-YAP pathway plays essential roles during heart development and heart regeneration (*4*). While the deletion of YAP in embryonic heart impairs cardiomyocyte division, the expression of constitutively active YAP promotes cardiomyocyte proliferation (*5*, *9*, *33*, *34*). We now show that MMB is required for the ability of YAP to induce cell division in this tissue (Figure 8F). First, we used a mouse model in which cell cycle genes were induced in embryonic hearts by activation of YAP through the knockout of the Hippo pathway protein *Sav1*. These genes were not induced when the MuvB subunit *Lin9* was deleted together with *Sav1*. Proliferation of cardiomyocytes is also dependent on both MMB and YAP since deletion of *Lin9* abolishes the enhanced proliferation induced by deletion of *Sav1*. This requirement is not limited to the embryonic heart but the same dependency on *Lin9* for cardiomyocyte proliferation was also observed in postnatal hearts using *Lin9* / *Sav1* double knockout mice expressing the Cre transgene from the alpha-MHC promoter. Further evidence for this dependency on LIN9 comes from experiments using cultured embryonal or postnatal cardiomyocytes with a conditional *Lin9* allele and a hormone inducible Cre recombinase. Adenoviral expression of activated YAP was only able to induce mitosis when *Lin9* was present but not after Cre-mediated deletion of *Lin9*. Although it is possible that YAP and MMB could independently induce a similar set of genes required for cell cycle regulation, our findings that B-MYB and YAP directly interact and that YAP induced cardiomyocyte proliferation depends on its WW domains supports the notion that the synergy is a result of the interaction between YAP and MMB. Furthermore, cardiomyocyte mitosis is inhibited by MY-COMP, a fragment of B-MYB that contains the YAP binding domain of B-MYB. When fused to a nuclear localization signal and expressed in cells, MY-COMP interferes with the association YAP to B-MYB. Genome binding studies in embryonic hearts demonstrated that LIN9 binds to the promoters of cell cycle genes activated by YAP, indicating that these genes are direct targets of MMB. By ChIP-seq, YAP does not colocalize with LIN9 to the promoters of cell cycle genes but instead binds to enhancers, consistent with recent data from human cancer cell lines where YAP also predominantly binds to distant sites (*24*) (*25*, *26*). This suggests that YAP interacts with MMB-regulated promoters through chromatin looping, resulting in the activation of a subset of cell cycle genes, similar to what we have recently described for human MCF10A cells (*14*). Given that YAP not only interacts with MMB but that the MMB subunit B-MYB is also a transcriptional target of YAP, one can speculate that YAP and B-MYB form a positive feedback loop enabling expression of downstream cell cycle genes.

Remarkably, although LIN9 was required for cardiomyocyte cell cycle induction due to activated YAP, LIN9 was completely dispensable for homeostasis of the adult murine heart. This absence of a phenotype in adult cardiac-specific *Lin9* knockout mice was surprising, because LIN9 is a key subunit of the DREAM complex that is involved in repression of cell cycle genes in non-dividing cells (*15*). The observation that the genetic inactivation of *Lin9* in the postnatal heart is not associated with ectopic proliferation of cardiomyocytes suggests that loss of DREAM does not lead to sufficient de-repression of cell cycle genes to induce cardiomyocyte proliferation. It is therefore possible that DREAM does not permanently silence cell cycle genes, but keeps them in a poised state for re-activation by pro-proliferative signals, consistent with our finding that LIN9 remains bound to promoters in postnatal hearts. Also supporting this possibility is finding that cell cycle promoters, as opposed to promoters of genes that regulate cell-cell contacts and the cytoskeleton, are readily accessible in adult murine hearts before introduction of a YAP5SA transgene (*6*). Thus, MuvB complexes may have a dual role in cardiomyocyte proliferation, contributing to the inactive, but primed state of cell cycle genes as well as to their YAP-mediated activation. It will be important to investigate whether epigenetic changes at MuvB bound promoters contribute to the loss of the regeneration capacity of adult hearts (*35*).

YAP not only plays a role in cardiac development but is also a potent oncogene in different cancers. Previous findings have implicated the YAP WW domain in oncogenic transformation (*36*). Targeting WW domain mediated interactions by MY-COMP, or by more potent inhibitors based on MY-COMP, may prove a valid strategy to inhibit the oncogenic pro-proliferative functions of YAP. Such inhibitors could serve as selective therapeutic target in cancers with high levels of YAP.

## MATERIALS AND METHODS

### Mice

*Sav1*^*tm1.1Dupa*^/J mice were obtained from Jackson laboratories, in which LoxP sites flank exon 3 of the *Sav1* gene (*23*). We have previously descripted *Lin9*^*f*l^ mice, in which LoxP sites flank exon 7 of the *Lin9* gene (*37*). Cardiomyocyte-specific deletion of *Sav1* and *Lin9* was achieved by crossing mice to Nkx2.5-Cre mice (*38*). Postnatal cardiomyocyte-specific deletion of *Sav1* and *Lin9* was achieved by crossing mice to α-MHC-Cre mice (*27*). To obtain Lin9^fl/fl^CreER^T2^, *Lin9*^*fl*^ mice were crossed with a mouse line ubiquitously expressing CreER^T2^ transgene (*39*). Mice were maintained in a C57/Bl6 background.

### Cell culture

HEK293A (Thermo Fisher Scientific) cells, HeLa (ATCC® CCL-2™, female) cells and C2C12 (ATCC® CRL-1772™) cells were maintained in DMEM supplemented with 10% FCS (Thermo Fisher Scientific) and 1% penicillin/streptomycin (Thermo Fisher Scientific). Differentiation of C2C12 cells was induced by culturing in DMEM medium with 2 % horse serum (Sigma). Primary embryonic and postnatal cardiomyocytes were isolated by enzymatic digestion with the Pierce™ Primary cardiomyocyte isolation kit (Thermo Fisher Scientific). Cardiomyocytes were enriched by preplating on tissue-culture plastic to remove nonmyocytes. Cardiomyocytes were initially cultured for 48h on fibronectin-coated dishes with 5% horse serum (Sigma) and 1% penicillin/streptomycin (Thermo Fisher Scientific) and cardiomyocyte growth supplement at 37 °C and 5% CO2 to prevent proliferation of nonmyocytes. To induce the deletion of *Lin9*, conditional cardiomyocytes were treated with 100 nM 4-hydroxytamoxifen (Sigma) for 4 days. Cardiomyocytes were transduced with adenovirus (multiplicity of infection, 25) in serum-free medium for 24 h and cultured for additional 48 h.

### Adenovirus

Adenoviral constructs were constructed using the ViralPower adenoviral expression system (Thermo Fischer Scientific). Entry vectors containing YAPS127A, YAPS94AS127A and YAPS127AΔ155-263 were generated by PCR from p2xFLAG-CMV-YAPS127A (*40*). Entry vectors containing GFP, HA-NLS-B-MYB (2-241)-T2A-GFP and HA-NLS-B-MYB (2-241, ΔPPXY)-T2A-GFP were generated by PCR from pCDNA4/TO-HA-NLS-SV40-B-MYB (this work) and pSpCas9n(BB)-2A-GFP (PX461) (Addgene #48140). All entry vectors were subcloned into pENTR3C (Invitrogen) and then recombined into pAd/CMV/V5-DEST using LR clonase II (Thermo Fischer Scientific). The production of YAP mutant and LacZ adenoviruses was performed in HEK293A cells according to the manufacturer’s instructions. Adenoviruses were titered using the Adeno-X Rapid Titer Kit (TaKaRa).

### RT-qPCR

Total RNA was isolated using peqGOLD TriFast (Peqlab) according to the manufacturer’s instructions. RNA was transcribed using 100 units RevertAid reverse transcriptase (Thermo Fisher Scientific). Quantitative real–time PCR reagents were from Thermo Fisher Scientific and real-time PCR was performed using the Mx3000 (Stratagene) detection system. Primer sequences are listed in Supplementary Table 1. Expression differences were calculated as described before (*20*).

### PLA

PLA was performed using the Duolink In Situ Kit (Sigma) according to the manufacturer’s instructions. The following antibodies were used: LIN9 (Bethyl, A300-BL2981; 1:150), B-MYB phospho-T487 (ab76009; 1:100), YAP (Santa Cruz Biotechnology, sc-101199;1:200). For secondary staining, samples were incubated for 45 minutes with the following antibodies: cTnT (DSHB; 1:50) or α-HA (Sigma, 1:200). After washing several times with buffer A, samples were incubated with for 20 minutes with secondary antibody: α-IgG2a conjugated to Alexa 488 (Thermo Fisher Scientific) and Hoechst 33258 (Sigma). Pictures were taken with an inverted Leica DMI 6000B microscope equipped with a Prior Lumen 200 fluorescence light source and a Leica DFC350 FX digital camera.

### Plasmids

Expression vectors for truncated human FLAG-YAP constructs (aa 2-499, 2-287, 2-263, 2-154, 155-504, 264-504, 288-504 and Δ155-263) were constructed from pCMV-2xFLAG-YAP (*41*) by PCR. Expression vector for human GST-YAP constructs (aa 2-263, 155-263, 168-204, 231-263) were constructed by PCR in the pGEX-4-T2 vector. Truncated human HA-B-MYB constructs (aa 2-410, 2-331, 80-331, 80-410, 2-241, 2-79, 242-410) were generated from pCDNA4/TO-HA-B-MYB. Expression vector for truncated human 6xHis-B-MYB-2-241 and 2-241Δ198-201 (ΔPPXY) were constructed by PCR using appropriate oligonucleotides and were subcloned into pRSETA. Expression vectors containing HA-NLS-SV40-B-MYB-2-241, 2-241 N174A(ΔDNA), 2-241 Δ198-201(ΔPPXY) were obtained by PCR and were subcloned into pCDNA4/TO vector.

### Immunoblotting and immunoprecipitation

Whole protein extracts were obtained by lysing cells in TNN buffer (50mM Tris (pH 7.5), 120mM NaCl, 5mM EDTA, 0.5% NP40, 10mM Na4P_2_O_7_, 2mM Na_3_VO_4_, 100mM NaF, 1mM PMSF, 1mM DTT, 10mM β-glycerophosphate, protease inhibitor cocktail (Sigma)). Protein lysates or purified protein were separated by SDS-PAGE, transferred to a PVDF-membrane and detected by immunoblotting with the first and secondary antibodies: β-actin (Santa Cruz Biotechnology, sc-47778) 1:5000, B-MYB (clone LX015.1, (*42*) 1:5, anti-HA.11 (Covance, MMA-101P) 1:1000, anti-FLAG M2 (Sigma, F3165) 1:5000, anti-His (Sigma, H1029) 1:2000, Vinculin (Sigma, V9131) 1:10000, TOP2A (Santa Cruz Biotechnology, sc-365916) 1:1000, CDC20 (Santa Cruz Biotechnology, sc-5296) 1:500, YAP (Santa Cruz Biotechnology, sc-10199) 1:1000, p-YAP(S127A) (Cell Signaling Technology, 4911) 1:1000, LIN9 (Bethyl, A300-BL2981), NUSAP1 (Geert Carmeliet) 1:1000, CENPF (Abcam, ab-5) 1:1000, anti-mouse-HRP (GE healthcare, NXA931) 1:5000 and HRP Protein A (BD Biosciences, 610438) 1:5000. For immunoprecipitation of FLAG-tagged proteins, protein G dynabeads (Thermo Fisher Scientific) were first coupled with 1 µg FLAG-antibody (Sigma, F3165) and then immunoprecipitated with 1mg whole cell lysate. After five times of washing with TNN, proteins were separated by SDS-PAGE and detected by immunoblotting using the desired antibodies.

### Immunostaining

For immunostaining cardiomyocytes were seeded onto fibronectin coated coverslips. HeLa cells were seeded onto coverslips without fibronectin. Cells were fixed with 3% paraformaldehyde and 2% sucrose in PBS for 10 minutes at room temperature. Cells were permeabilized using 0.2% Triton X-100 (Sigma) in PBS for 5 minutes and blocked with 3% BSA in PBS-T (0.1% Triton X-100 in PBS) for 30 minutes. Primary antibodies were diluted in 3% BSA in PBS-T and incubated with the cells for 1 hour at room temperature. The following antibodies were used: TroponinT (CT3) (developmental studies hybridoma bank) 1:50, phospho-Histone H3 (Ser10) (Santa Cruz Biotechnology, sc-8656) 1:100, Ki-67 (SP6) (Thermo Scientific, RM-9106) 1:200, α-tubulin (B-5-1-2) (Santa Cruz Biotechnology, sc-23948) 1:150 and anti-HA (Sigma, H6908) 1:200. After three washing steps with PBS-T, secondary antibodies conjugated to Alexa 488 and 594 (Thermo Fisher Scientific) and Hoechst 33258 (Sigma) were diluted 1:500 in 3% BSA in PBS-T and incubated with the coverslips for 30 minutes at room temperature. Finally, slides were washed three times with PBS-T and mounted with Immu-Mount™ (Thermo Fisher Scientific). Pictures were taken with an inverted Leica DMI 6000B microscope equipped with a Prior Lumen 200 fluorescence light source and a Leica DFC350 FX digital camera.

### Histology, H&E staining and immunohistochemistry

Freshly dissected embryos and postnatal hearts were fixed with bouin’s fixative (picric acid (saturated; AppliChem), 10% (v/v) formaldehyde, 5% (v/v) (AppliChem) glacial acetic acid (Roth) embedded in paraffin and sectioned. Sections were deparaffinized, rehydrated and stained with hematoxylin and eosin or processed for immunostaining.

Following deparaffinization, antigen retrieval was performed by boiling samples in 10mM sodium citrate buffer (pH6.0) for 10 minutes in a microwave oven. After 30 minutes cool down, samples were blocked with 3% BSA in PBS-T (0.1% Tween-20 (AppliChem)) and incubated with the primary antibodies diluted in PBS-T over night at 4°C. The following antibodies were used: TroponinT (CT3) (developmental studies hybridoma bank) 1:50, phospho-Histone H3 (Ser10) (Santa Cruz Biotechnology, sc-8656) 1:100, Ki-67 (SP6) (Thermo Scientific, RM-9106) 1:200. After three washing steps with PBS-T, secondary antibodies conjugated to Alexa 488 and 594 (Thermo Fisher Scientific) and Hoechst 33258 (Sigma) were diluted 1:200 in 3% BSA in PBS-T and incubated with the coverslips for 2 hours at room temperature. Finally, slides were washed three times with PBS-T and mounted with Immu-Mount™ (Thermo Fisher Scientific). Pictures were taken with an inverted Leica DMI 6000B microscope equipped with a Prior Lumen 200 fluorescence light source and a Leica DFC350 FX digital camera.

### Flow cytometry

Cardiomyocytes were isolated and enriched with the Pierce™ Primary cardiomyocyte isolation kit (Thermo Fisher Scientific) according to the manufacturer’s instructions. Isolated and enriched cardiomyocytes were fixed with 4% PFA in PBS for 5 minutes. Detection was performed using a Beckman Coulter FC 500 cytomics and data were analyzed in CXP analysis 2.2 software. Gating and compensation were based on fluorophore-negative controls. 10.000 cells were analyzed per genotype.

### RNA-Seq

For whole transcriptome analysis, total RNA was isolated in triplicates from ventricular heart tissue with the desired genotype. DNA libraries were generated using 1µg RNA with the magnetic mRNA isolation module and NEBNext Ultra II RNA Library Prep Kit for Illumina (New England Biolabs). DNA library was amplified by 7 PCR cycles and quality was analyzed using the fragment analyzer (Advanced Analytical). Libraries were sequenced on the NextSeq 500 platform (Illumina).

### ChIP-Seq

Chromatin was isolated from ventricular heart tissue of E16.5 or P10 hearts. In brief, minced tissue was fixed with 1% formaldehyde for 20 minutes at RT. The reaction was stopped by adding 125mM glycine for additional 5 minutes. Tissue was incubated in lysis buffer (50mM Tris-HCl pH8, 10mM EDTA, 1% SDS) for 1 hour at 4°C. Samples were sonicated for 1 minute at 25% amplitude (10s ON / 30s OFF) at 4°C, insoluble material was removed by centrifugation and chromatin was fragmented to an approximate size of 150 to 300bp by additional sonicating for 10 minutes at 25% amplitude (10 seconds On/ 30 seconds OFF) at 4°C using a Branson sonifier. Afterwards chromatin was diluted ten times in ChIP dilution buffer (50mM Tris-HCl pH8, 0.167M NaCl, 1.1% Triton X-100, 0.11% sodium deoxycholate). For immunoprecipitation, 9µg of the antibody was coupled to protein G dynabeads (Thermo Fisher Scientific) for 6 hours at 4°C and then incubated with fragmented chromatin over night at 4°C. Beads were washed in total twelve times with wash buffer I (50mM Tris-HCl pH8, 0.15M NaCl, 1mM EDTA, 0.1% SDS, 1% Triton X-100, 0.1% sodium deoxycholate), wash buffer II (50mM Tris-HCl pH8, 0.5M NaCl, 1mM EDTA, 0.1% SDS, 1% Triton X-100, 0.1% sodium deoxycholate), wash buffer III (50mM Tris-HCl pH8, 0.5M LiCl_2_, 1mM EDTA, 1% Nonidet P-40, 0.7% sodium deoxycholate) and wash buffer IV (10mM Tris-HCl pH8, 1mM EDTA). 1mM PMSF and protease inhibitor cocktail were added freshly to all buffers. After washing chromatin was eluted in (10mM Tris-HCl pH8, 0.3M NaCl, 5mM EDTA, 0.5% SDS, 10µg/ml RNAseA) and crosslink was reversed at 65°C over night. Proteins were digested by adding 200µg/ml proteinase K at 55°C for 2 hours. DNA was purified using the QIAquick PCR Purification Kit (QIAGEN) and eluted in 50 µl EB buffer. Purified ChIP-DNA was quantified using the Quant-iT PicoGreen dsDNA Assay Kit (Thermo Fisher Scientific). DNA libraries were generated using 10ng purified ChIP-DNA and the NEBNext®Ultra IIDNALibrary Prep Kit for Illumina® (New England Biolabs) according to the manufacturer’s instructions. DNA library was amplified by 12-15 PCR cycles and quality was analyzed using the fragment analyzer (Advanced Analytical). Libraries were sequenced on the NextSeq 500 platform (Illumina). The following antibodies were used for ChIP-Seq: LIN9 (Bethyl, A300-BL2981), YAP (NB110-58358) and IgG from rabbit serum (Sigma, I5006).

### GST-pulldown

Recombinant GST, GST-TEAD-WW1/2 (TEAD-binding domain and WW domain 1 and 2 of YAP fused to GST), GST-WW1/2-, GST-WW1, GST-WW2 or His-B-MYB(2-241) (amino acids 2-241 fused to 6 x his) were expressed in BL21 cells and purified on glutathione-linked sepharose beads (GE healthcare) or Ni-NTA-Agarose (GE healthcare, His-B-MYB). Lysates of HeLa cells expressing HA-tagged B-MYB constructs were incubated with 5 mg immobilized GST or GST-WW1/2-YAP for 3 hours at 4°C. Beads were washed six times with TNN buffer, resuspended in SDS protein sample buffer, boiled for 5 min, separated on a 10% SDS-PAGE gel, blotted and analyzed via immunoblotting.

### Bioinformatic Analysis

After sequencing, bases were called using Illuminas GenerateFASTQ v1.1.0.64 software and sequencing quality was controlled with the FastQC script. For RNA-seq, reads were mapped with TopHat v2.1.0 (*43*) and BOWTIE2 (*44*) with default parameters to the murine genome (mm10). Samples were normalized to the sample with the smallest number of mappable reads and a read count matrix for each Ensembl gene (GRCm38.p6) was generated with the *summarizeOverlaps* function in the R package {GenomicFeatures}. Before differential gene expression analysis, non- or weakly expressed genes were removed using the threshold: mean read count per gene over all samples >1. Differentially expressed genes were called with *EdgeR* and p-values were adjusted for multiple-testing with the Benjamini-Höchberg procedure (FDR: false discovery rate). Gene set enrichment analyses were performed with signal2noise metric, 1000 permutations and a combined gene set database comprising Hallmark and C2 gene sets.

For ChIP-seq, sequenced reads were mapped to the *Mus musculus* genome mm10 with BOWTIE v1.2 (*44*) with default parameters and subsequently normalized to the sample with the smallest number of sequenced reads. Peaks were called with MACS14 (*45*) with maximal 3 duplicates, a p-value cut-off of 1e-5 and the input sample as control. Resulting peaks were annotated to the next transcriptional start of Ensembl genes with the *closestBed* function from the bedtools suite v2.26.0 (*46*). Overlapping peaks were identified with *bedtools intersect* and a minimal overlap of 1bp. LIN9 occupancy was calculated in a window of +/-1kb around TSSs with *bedtools coverage* function. Density matrices were generated with deeptools v2.3.5. (*47*) *computeMatrix* function at a resolution of 1bp and subsequently used for plotting heat maps with *plotHeatmap*. Mapped ChIP-seq data for histone marks were taken from the ENCODE portal (*48*) (https://www.encodeproject.org/) with the following identifiers: ENCFF056JGV, ENCFF295HNV, ENCFF642EEK, ENCFF687BWU. Promoters, enhancers and super-enhancers were defined as described previously (*14*).

In box plots, the central line reflects the median, the borders of the boxes indicate the first and third quartile and whiskers extend to 1.5 of the interquartile range. Outliers are shown as dots. P-values were calculated with a two-tailed Wilcoxon rank sum test (unpaired samples) or Wilcoxons signed-rank test (paired samples). ChIP- and RNA-sequencing datasets are available at the NCBI’s Gene Expression Omnibus (*49*) under the accession number GEO: GSE137132.

## ACKNOWLEDGEMENTS

We thank Stefanie Hauser and Grit Weinstock for critical reading of the manuscript. This work was supported by grants from the Deutsche Krebshilfe (70113138) and Deutsche Forschungsgemeinschaft, DFG (GA575/9-1) towards SG.

## AUTHOR CONTRIBUTIONS

S.G. and M.G. planned the study and designed the experiments. M.G., L.H., M.S., K.M.W. and S.P. conducted the experiments. M.G. and S.G. analyzed the data. S.W. performed the bioinformatic analyses. C.P.A. performed next-generation sequencing. S.G. and M.G. wrote the manuscript.

