## Supplemental Material for "Interaction of YAP with the Myb-MuvB (MMB) complex defines a transcriptional program to promote the proliferation of cardiomyocytes"

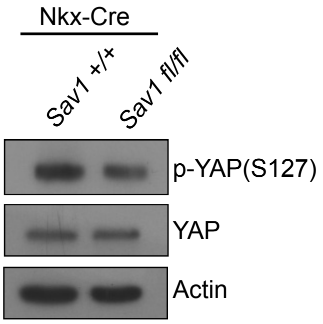


**Supplementary Figure S1: Proliferation of embryonic cardiomyocytes following Hippo inactivation depends on LIN9.** Expression of YAP and levels of YAP phosphorylated on S127 (p-YAP) in the hearts of Nkx2.5-Cre;*Sav1*$^{+/+}$ and Nkx2.5-Cre;*Sav1*$^{fl/fl}$ mice was determined by immunoblotting. Actin served as control.

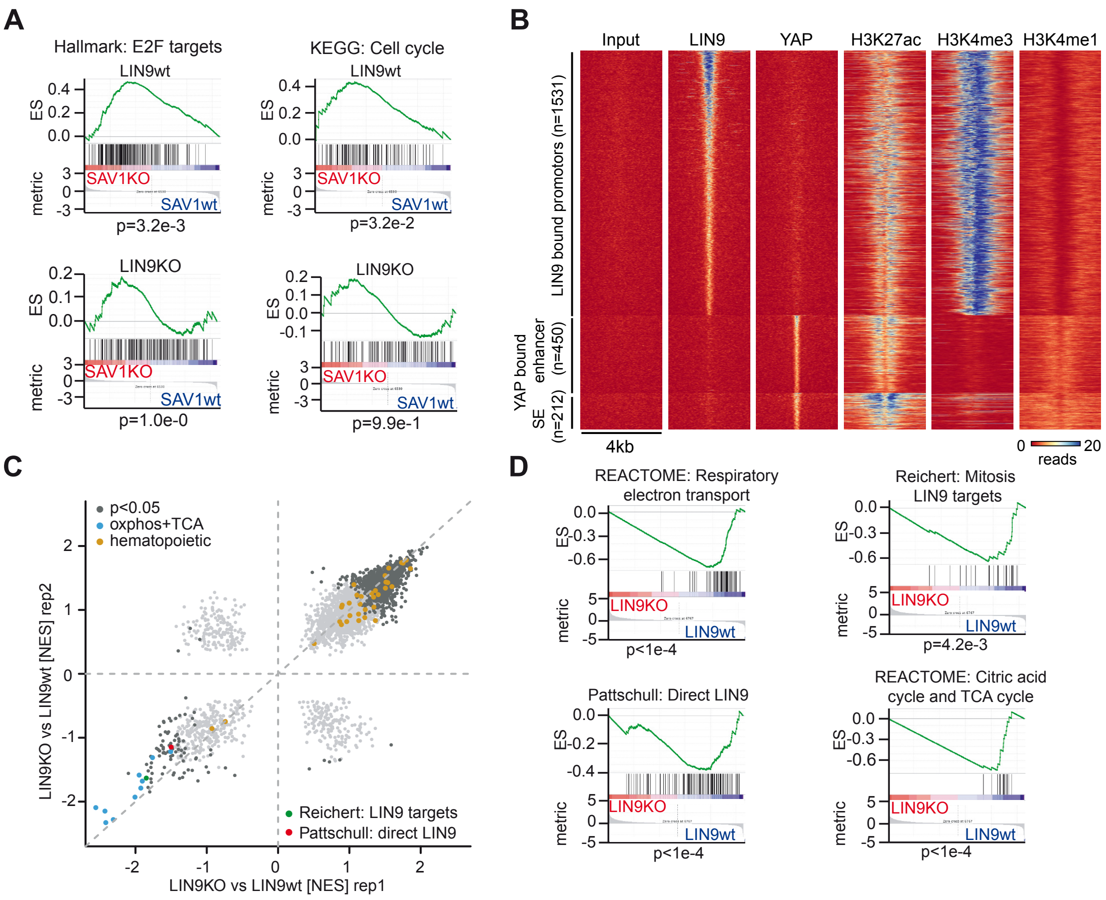


**Supplementary Figure S2: Cell cycle genes upregulated in *Sav1* knockout hearts are direct targets of LIN9** A) Representative gene sets from the analysis in Figure 5A. p-values were calculated using a permutation test with 1000 permutations. „Signal2Noise" was used as a metric to rank genes. ES: enrichment score. B) Heat map documenting binding of LIN9 and YAP at LIN9 peaks in promoters or at YAP peaks in enhancers and superenhancers in E16.5 heart ventricles. Read density is plotted in a window of +/-2kb around the peak at a resolution of 2bp. Data for histone modifications are taken from ENCODE. C) GSEA comparing expression differences in Nkx-Cre; Lin9 $^{fl/fl}$ (LIN9 KO) and Nkx-Cre; Lin9$^{fl/+}$ (LIN9 wt) heart ventricles from E13.5 mice in two biological replicates (each done in triplicate). The C2 MSigDB was spiked with the Hallmark gene sets and a set of LIN9 direct targets genes from (Pattschull *et al*, 2019). Gene sets related to respiration/TCA cycle ("oxphos") and hematopoietic cells are highlighted in blue and orange, respectively. NES: normalized enrichment score. D) Representative gene sets from the analysis in C. p-values were calculated using a permutation test with 1000 permutations. „Signal2Noise" was used as a metric to rank genes. ES: enrichment score.

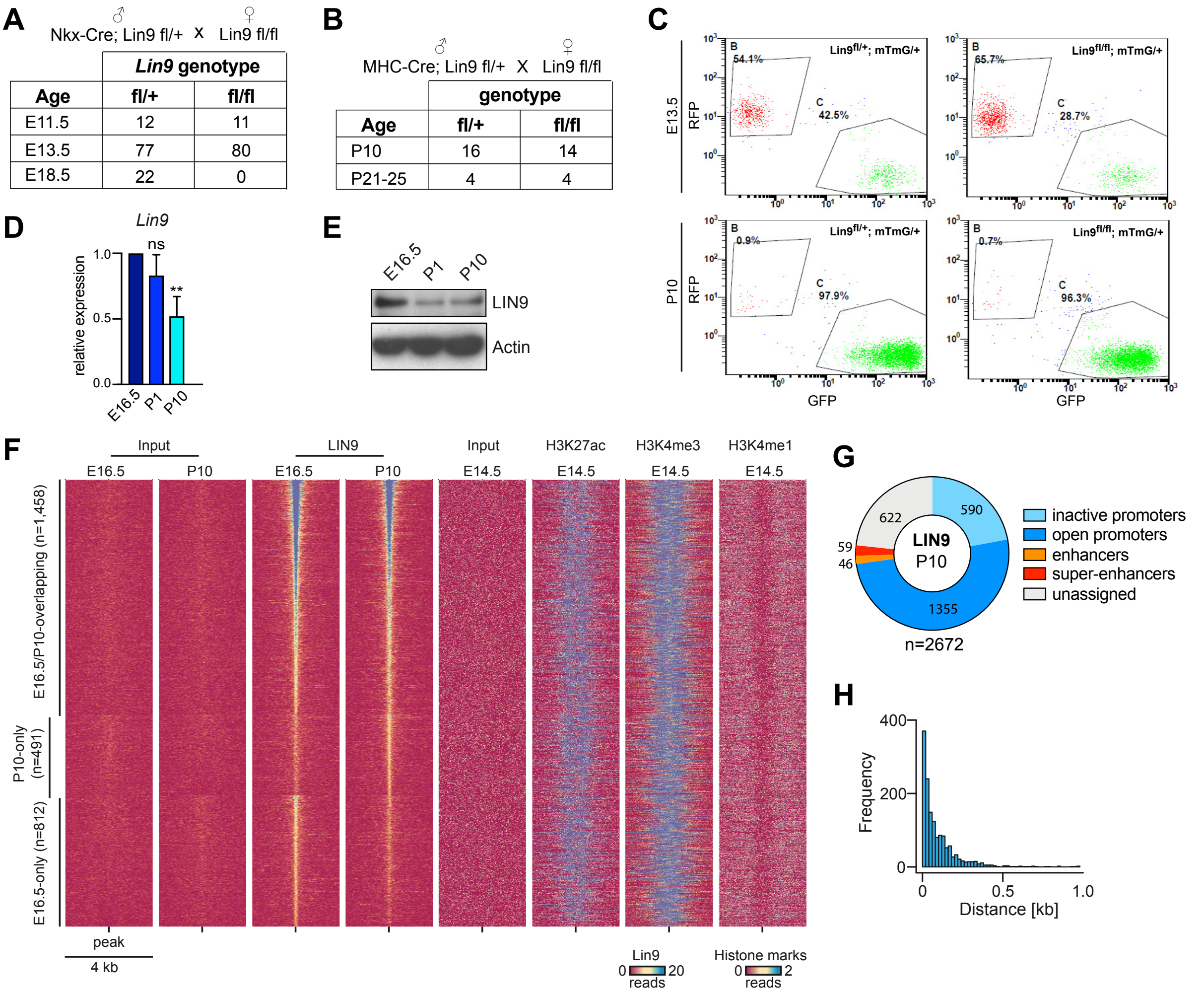


**Supplementary Figure S3: LIN9 is required for cardiomyocyte proliferation in Hippo-deficient, postnatal hearts** A) Embryonic lethality of *Nkx2.5*$^{Cre}$*; Lin9*$^{fl/fl}$ mice. Breeding scheme and resulting genotypes. Result of the genotyping of live embryos at the indicated developmental time points. B) Viability of alpha-MHC-Cre*; Lin9*$^{fl/fl}$ mice Breeding scheme and resulting genotypes. Number of mice with the indicated genotypes at P10 and P21-25. C) Example for a FACS analysis of mTomato and mEGFP positive cardiomyocytes derived from hearts of E13.5 and P10 hearts with the indicated genotypes. See Figure 3G. D) The expression of *Lin9* relative to *Actin and Hprt* was investigated in E16.5, P1 and P10 hearts by RT-qPCR. n= 3 independent replicates. E) The expression of LIN9 in lysates prepared from hearts at the different developmental stages was investigated by immunoblotting. β-actin served as a loading control. F) Heat map documenting binding of LIN9 at LIN9 peaks in promoters called in E16.5 or P10 cardiomyocytes or overlapping peaks. Read density is plotted in a window of +/-2kb around the peak at a resolution of 2bp. Data for histone modifications are taken from ENCODE (GSE31039). G) Plot illustrating the genomic localization of LIN9 in postnatal (P10) heart ventricles as determined by ChIP-seq. H) Histogram showing the absolute distance between overlapping LIN9 peaks called in E16.5 and P10 heart ventricles located in promoters (n=1,458) at a resolution of 20 bp.

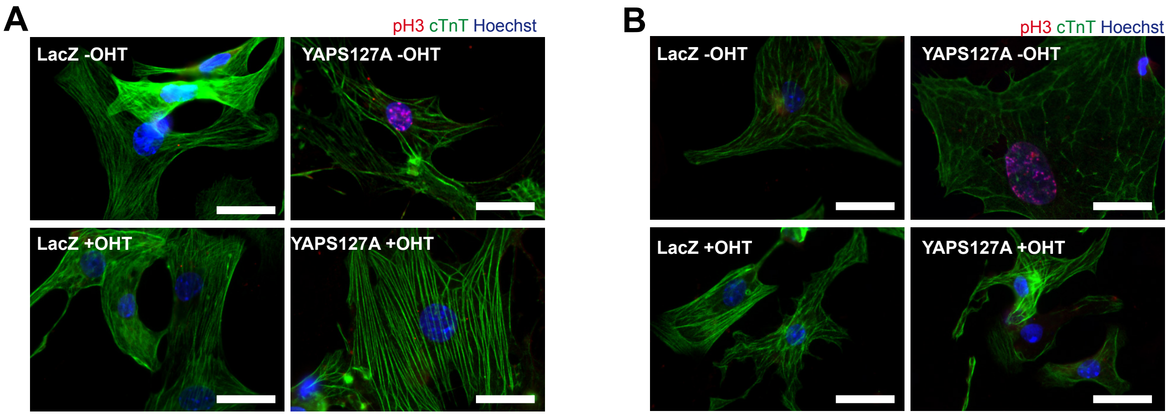


**Supplementary Figure S4: Cardiomyocyte proliferation by activated YAP requires MMB**

A) -B) Embryonal (E14.5) *Lin9*$^{fl/fl}$;CreER cardiomyocytes (A) or postnatal (P1) cardiomyocytes (B) were transduced with Ade-LacZ or with Ade-YAP(S127A) and treated with or without 4-OHT. The fraction of pH3-positive cardiomyocytes was quantified by staining for pH3 (red). Example microphotograph of the experiments shown in Figure 4C and D.

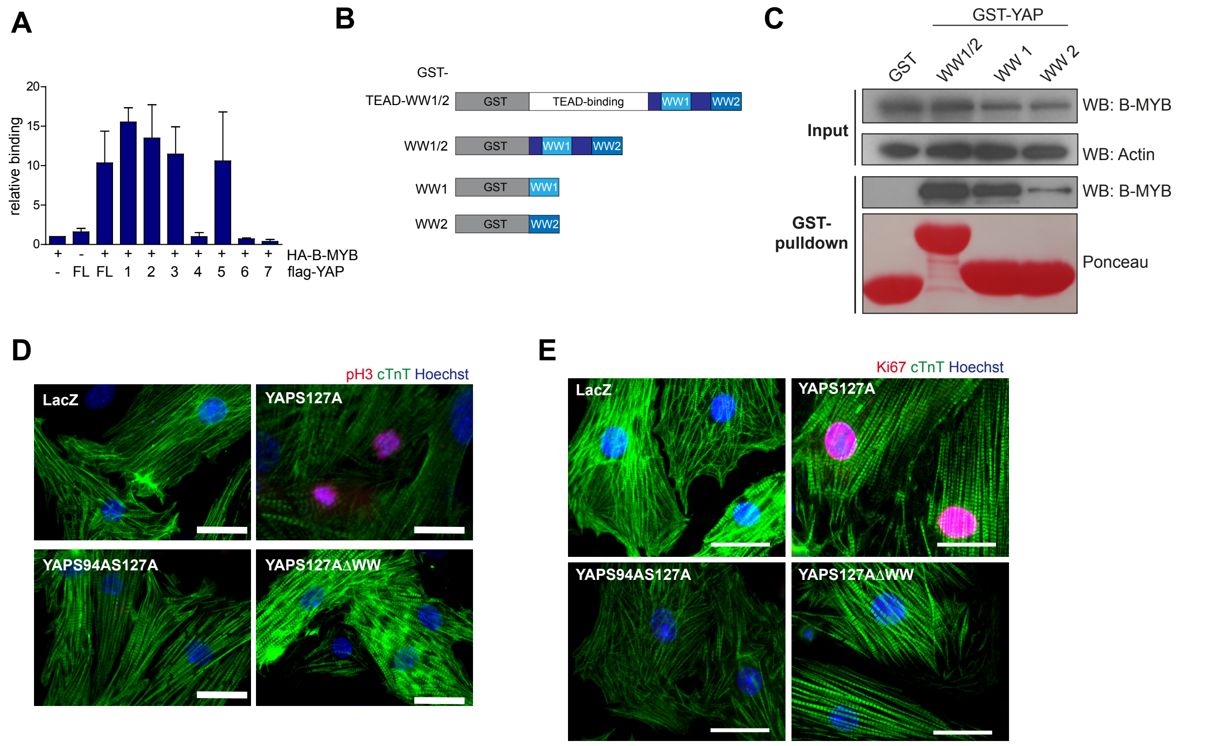


**Supplementary Figure S5: The WW domains of YAP mediate the interaction with B-MYB and are required to induce cardiomyocyte proliferation** A) Densiometric quantification of binding data shown in Figure 6B using ImageJ. Binding is relative to HA-B-MYB control cells. n=3 biological replicates. B) Scheme of the GST fusion constructs used in pulldown experiments in Figure 6D and S5C. C) Pulldown experiments of the indicated GST fusion proteins with HA-B-MYB. Bound B-MYB was detected by immunoblotting with an HA-antibody. Input: 3 % of the lysate used for the pulldown was loaded onto the gel. Actin served as a control. Ponceau staining was used to detect the recombinant GST-proteins. D) and E) Cardiomyocytes were transduced with Ade-LacZ, Ade-YAP(S127A), and Ade-YAP(S127A/S94A) or with Ade-YAP(S127A/ΔWW). The fraction of pH3-positive positive (D) and Ki67 (E) cardiomyocytes was determined by immunostaining. Example microphotographs of the experiment shown in Figure 6E and F.

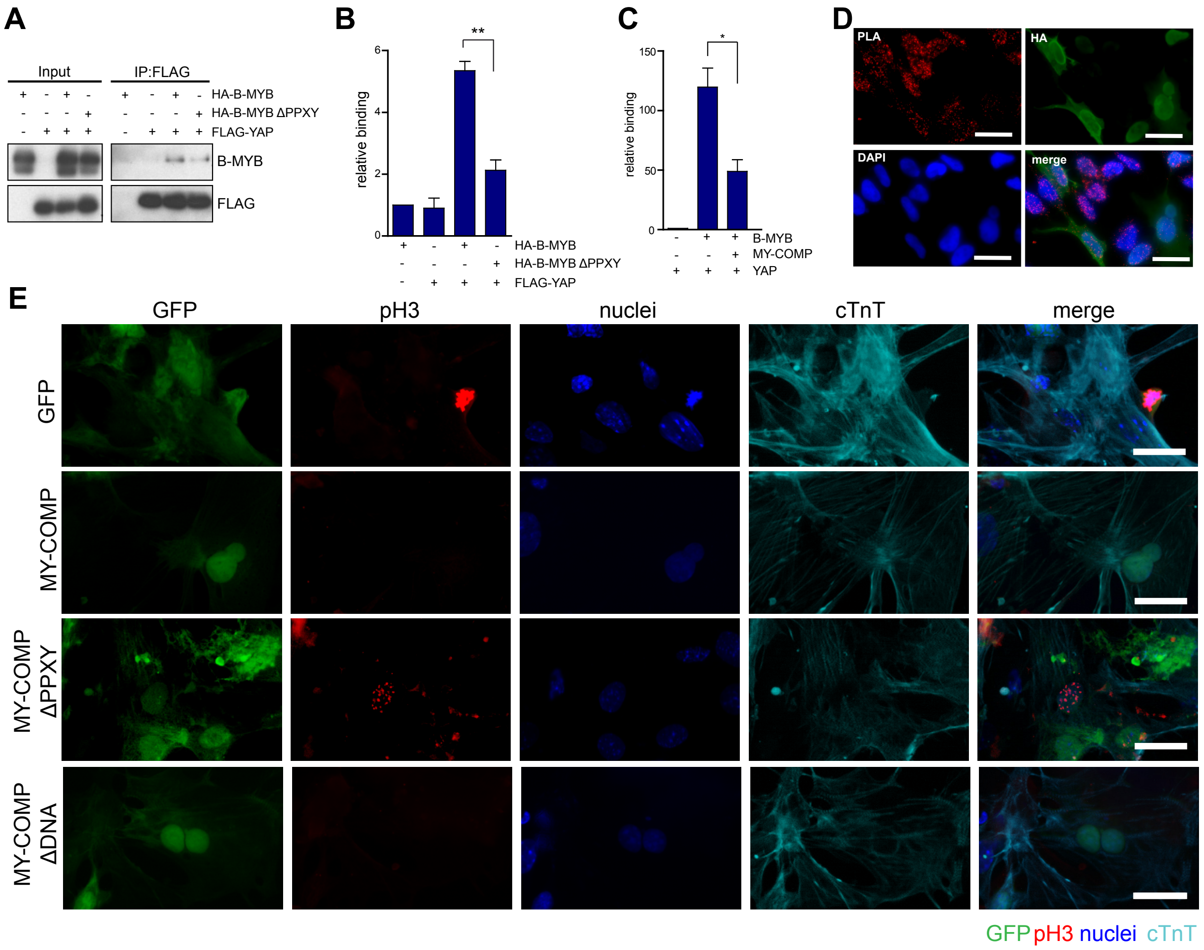


**Supplemental Figure S6: Disrupting the association between B-MYB and YAP by MY-COMP inhibits cardiomyocyte proliferation** A) Co-immunoprecipitation experiments of the indicated B-MYB constructs with flag-YAP. Lysates of HeLa cells expressing flag-YAP and HA-B-MYB (wt and ΔPPXY) were immunoprecipitated with flag-antiserum and immunoblotted with an anti-HA-antibody and anti-flag antibody. 3 % percent of total lysate was immunoblotted (Input). B) Densiometric quantification of the binding of HA-MYB and HA-B-MYBΔPPXY to flag-YAP using Image J. Binding is relative to HA-B-MYB control cells. C) Densiometric quantification of the binding of HA-B-MYB to flag-YAP in presence and absence of MY-COMP using Image J. Binding is relative to FLAG-YAP control cells. N=3 biological replicates. D) Proximity ligation assay (PLA) of endogenous YAP and B-MYB upon transfection of MY-COMP. Cell expressing MY-COMP were identified by HA staining (green). Example microphotographs of the experiments shown in Figure 7F. E) Embryonal cardiomyocytes were infected with adenoviruses expressing GFP, HA-NLS-B-MYB(2-241), with HA-NLS-B-MYB(2-241,ΔPPXY) or with HA-NLS-B-MYB(2-241,ΔDNA) each coupled to GFP through a T2A self-cleaving peptide. Mitotic cells were quantified by staining for phospho-H3 (red). B), C): Error bars indicate SEMs. Student's t-test. $*=p<0.05$, $**=p<0.01$.

**Supplementary Table 1**

### RT-qPCR primers for murine genes

| Gene | fw primer | bw primer |
|---|---|---|
| *Actin* | GCTACAGCTTCACCACCACA | AAGGAAGGCTGGAAAAGAGC |
| *Anln* | ACAATCCAAGGACAAACTTGC | GCGTTCCAGGAAAGGCTTA |
| *Aspm* | GATGGAGGCCGAGAGAGG | CAGCTTCCACTTTGGATAAGTATTTC |
| *Axl* | TGCATGAAGGAATTTGACCA | ATCTGAGTGGGCAGGAACAC |
| *Birc5* | CCCGATGACAACCCGATA | CATCTGCTTGACAGTGAGG |
| *Casc5* | CACTCAGTTTTCAGTCTCTGTTAGATG | TGAAAAATAAGCTTGTGAACTAAAAGG |
| *Ccna2* | CTTGGCTGCACCAACAGTAA | CAAACTCAGTTCTCCCAAAAACA |
| *Cenpf* | AGCAAGTCAAGCATTTGCAC | GCTGCTTCACTGATGTGACC |
| *Ctgf* | TGACCTGGAGGAAAACATTAAGA | AGCCCTGTATGTCTTCACACTG |
| *Cyr61* | CAGCACCTCGAGAGAAGGAC | GGTCAAGTGGAGAAGGGTGA |
| *Ect2* | TGCTCTGCTTCACTGGATTC | ATGATGAACCAACGTCACCA |
| *Foxm1* | ACTTTAAGCACATTGCCAAGC | GGAGAGAAAGGTTGTGACGAA |
| *Gas2l3* | GCAGCCTGCAATCCAAGT | AGGGGACACCTGGGACTTA |
| *Hprt* | TCCTCCTCAGACCGCTTTT | CCTGGTTCATCATCGCTAATC |
| *Kif23a* | CTGTTGCCGTTGAAATGAGA | GGCTGTCAGTTCAAGGTTTCTT |
| *Lin37* | GTACCCCCGATGATGAACCT | CGTTGCATGTTGCGATAGAT |
| *Lin52* | GGTACGAGGCCTACAGAACCT | TCCCCTTGTCATCTCTCTGG |
| *Lin9* | TTGGGACTCACACCATTCCT | GAAGGCCGCTGTTTTTGTC |
| *Mybl2* | TTAAATGGACCCACGAGGAG | TTCCAGTCTTGCTGTCCAAA |
| *Nusap1* | TCTAAACTTGGGAACAATAAAAGGA | TGGATTCCATTTTCTTAAAACGA |
| *Prc1* | TGAGTCTATCACATGCCTGC | GCTCCTCTGGAATCCCAATT |
| *Sav1* | TCACACCAATAAAAGGGCTCA | CTTTCAGTTTGTTGTGGCTGA |
| *Top2a* | CAAAAGAGTCATCCCCCAAG | GGGGTACCCTCAACGTTTTC |
